# Anoikis resistance and metastasis of ovarian cancer can be overcome by CDK8/19 Mediator kinase inhibition

**DOI:** 10.1101/2023.12.04.569970

**Authors:** Mehri Monavarian, Resha Rajkarnikar, Emily Faith Page, Asha Kumari, Liz Quintero Macias, Felipe Massicano, Nam Y Lee, Sarthak Sahoo, Nadine Hempel, Mohit Kumar Jolly, Lara Ianov, Elizabeth Worthey, Abhyudai Singh, Igor B. Roninson, Eugenia V Broude, Mengqian Chen, Karthikeyan Mythreye

## Abstract

Anoikis resistance or evasion of cell death triggered by matrix detachment is a hallmark of cancer cell survival and metastasis. We show that repeated exposure to suspension stress followed by recovery under attached conditions leads to development of anoikis resistance. The acquisition of anoikis resistance is associated with enhanced invasion, chemoresistance, and immune evasion in vitro and distant metastasis in vivo. This acquired anoikis resistance is not genetic, persisting for a finite duration without detachment stress, but is sensitive to CDK8/19 Mediator kinase inhibition that can also reverse anoikis resistance. Transcriptomic analysis reveals that CDK8/19 kinase inhibition induces bidirectional transcriptional changes in both sensitive and resistant cells, disrupting the balanced reprogramming required for anoikis adaptation and resistance by reversing some resistance associated pathways and enhancing others. Both anoikis resistance and in vivo metastatic growth of ovarian cancers are sensitive to CDK8/19 inhibition, thereby providing a therapeutic opportunity to both prevent and suppress ovarian cancer metastasis.

## INTRODUCTION

Metastasis causes the majority of cancer-related deaths, yet metastatic cancers remain largely incurable. A critical step in metastasis is the ability of tumor cells to survive upon detachment from the extracellular matrix and primary tumor site, enabling their dissemination and circulation to distant sites. [1–3] Despite our understanding of these metastatic features, their therapeutic targeting remains underdeveloped.

Ovarian cancers (OC) comprising several subtypes, are among the most devastating of gynecological cancers (fifth in cancer deaths among women) and archetypal examples of cancers that leverage the metastatic hallmark of anoikis resistance for both transcoelomic/intraperitoneal and distant metastasis. [4–8] As malignant ascites accumulates, tumor cells must survive in suspension and evade cell death. [9–12] These cells can then colonize new peritoneal and mucosal surfaces, or return to primary tumor sites, as shown by clonal analysis. [13] The precise mechanisms by which cells acquire such anoikis resistance remain a subject of intense investigation. Suspension culture studies have been used extensively to understand such mechanisms, however most of them focus on single time points or long-term suspension culture studies. Despite these limitations, tumor intrinsic signals and pathways that change the ability of cells to undergo cell death upon matrix detachment [14–18] have been identified. These anoikis resistance mechanisms include but are not limited to transcriptional upregulation of critical survival genes, repression of pro-apoptotic genes, and transient expression changes in genes associated with antioxidant defense. Changes in the genes and pathways have also been associated with tumor growth and progression in ovarian and other cancers [19–23]. Coordinated regulation of specific reprogramming processes such as epithelial to mesenchymal transition (EMT) [24, 25], cadherin and integrin switching [26] by oncogenic pathways such as Ras/Erk, PI3k/AKT, Rho, MYC and TGF-β pathways also impact intraperitoneal (i.p.) OC survival and growth under anchorage independence. [27] However, only a few studies [28, 29] directly compare anoikis sensitive and resistant models to understand how anoikis resistance can be reached or prevented and the mechanisms underlying this process.

Given the significance of transcriptional changes to overall cancer progression, selective and potent inhibitors of transcription associated kinases [30] have emerged and are currently being evaluated in the clinic. Among these, CDK8 and CDK19 are of particular interest as they regulate gene expression programs in response to various cellular stresses and have shown promise in preventing adaptive resistance in other contexts. [31] CDK8 and CDK19 are closely related kinases associated with the transcriptional Mediator complex that both positively and negatively regulate transcription [32–34], and their inhibition affects different events associated with transcriptional reprogramming including EMT [24], cell differentiation [35] and gene expression changes in response to various signals and stressors. [32, 33] Notably, CDK8/19 inhibitors have reached clinical trials for solid tumors and leukemias (clinicaltrials.gov NCT03065010, NCT04021368, NCT05052255, NCT05300438) with utility specifically for gynecological cancers under examination. [36]

In this study we describe a model system that tests the effects of repeated exposures to detachment stress followed by attached re-growth to mimic potential in vivo scenarios. We delineate the impact of such repeat exposures to detachment stress on the development of adaptation to anoikis stress in different OC models. Our phenotypic and transcriptomic characterization of such anoikis resistant cells reveal non-genetic, transcriptional reprogramming, concomitant with a more aggressive phenotype in vitro and in vivo. We further show that both anoikis resistance and i.p. growth of OC can be suppressed by specific inhibition of CDK8/19 Mediator kinases, which can also reverse such acquired anoikis resistance. Further, transcriptomic analysis of the effects of CDK8/19 Mediator kinase inhibition reveal positive and negative changes to both core and stress associated transcriptional responses that rebalance the transcriptional response resulting from anoikis resistance. Our findings define a novel therapeutic strategy for counteracting anoikis resistance and metastasis by specific targeting of CDK8/19-regulated transcriptional reprogramming.

## RESULTS

### Attachment-detachment cycles confer anoikis sensitive ovarian cancer cells with resistance to cell death in suspension

To study how cells develop the ability to evade cell death upon loss of attachment (anoikis resistance), we first screened a broad panel of tumor cell lines that span a spectrum of commonly used OC cell line models, a pancreatic cancer cell line (PANC1), prostate cancer cell line (PC3), a primary population (non-immortalized) of tumor cells derived from the ascitic fluid of an OC patient (EOC15) and three previously established non-oncogenic immortalized ovarian surface and fallopian tube epithelial cell lines (IOSE144, FT282 and P201 and P210). [23][20, 37] The goal was to systematically define the anoikis sensitivity spectrum and identify cell lines that exhibited sensitivity to matrix detachment/suspension stress. Anoikis was measured by the percentage of live cells after plating cell lines in poly Hema coated ultra-low attachment conditions in their respective growth media for a fixed time of 24 hrs. All cell lines were plated at the same cell density in their growth media to minimize stress from other variables. The percent of live cells in suspension relative to the initial plating numbers hereby referred to as cell viability ranged from 36.1 % for IOSE144 to 125.2% for OVCAR5 (Figure 1A). Models exhibiting less than 100% viability when plated into suspension at 24 hrs were designated as ’anoikis sensitive’ (AnS) and models exhibiting 100% or more live cells plated into suspension at 24 hrs as intrinsically ‘anoikis resistant’ (AnR). A subset of cell lines that exhibited resistance at 24 hrs were further assessed for changes in viability for up to 72 hrs in suspension. HEYA8, PANC1 and TOV21G cells retained their live cell percentages and anoikis resistance for up to 72 hrs, whereas OVCA420 cells increased their cell death after 72 hrs in suspension (Supplemental Figure 1A). Hence, 72 hrs. was used as an anoikis sensitivity time point for OVCA420 alone in all subsequent experiments. EOC15, (non-immortalized ascites derived epithelial [23]) that had been maintained in attachment (Supplemental Figure 1B) also exhibited measurable cell death in suspension at 24 hrs. (Figure 1A).

**Figure 1.**
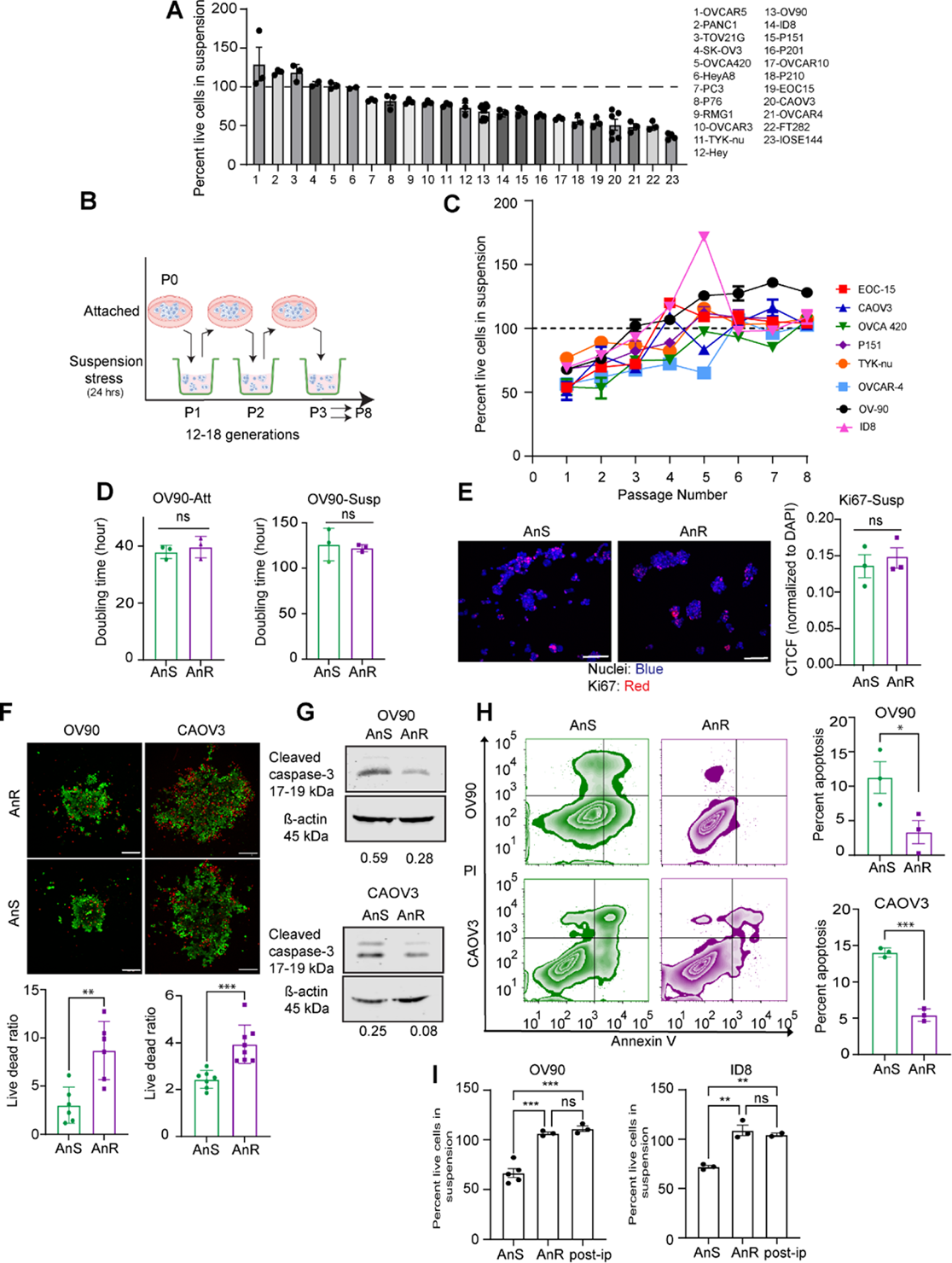
Development of anoikis resistance upon cyclic exposure to matrix detachment stress. **(A)** Percent live cells relative to initial plating number for the indicated cell lines after 24 hours in suspension assessed by a Trypan blue exclusion assay. (n≥3 for all cell lines). Mean ± SEM **(B)** Schematic of the cycles of attached growth followed by suspension culture conditions for testing development of resistance to repeated exposure to matrix detachment. **(C)** Percent live cells in suspension of indicated anoikis sensitive (AnS) cells measured by Trypan blue staining after 24 hours in suspension following cyclic gain and loss of attachment as in B. (n = 3-12 biological replicates per cell line). **(D)** Doubling time of OV90 cells either in attached (Att) growth conditions or in suspension (Susp) culture calculated over a 7 day or 10-day period respectively using an SRB assay. (Parental/AnS: standard culture conditions or AnR: anoikis resistant cells from between passages P7-P9 from C (n =2-3 independent biological trials with n =3 technical replicates). Mean ± SEM, ns p > 0.05, unpaired t-test. **(E)** Representative confocal images (left) and image quantitation (right) of the corrected total cell fluorescence (CTCF) of Ki67 (red) normalized to CTCF of DAPI of OV90 AnS and AnR derivative cells cytospun after 24 hours in suspension. ns p > 0.05, unpaired t-test, Mean ± SEM. Scale bar: 100 µm. (n = 3 biological trials, each trial includes quantitation of four 20X fields) **(F)** Live dead confocal images (above) and image quantiation (below) of indicated AnS and AnR derivative cells cultured for 24 hours in ultra-low attachment plates. Scale bar: 200 µm. Live dead cell ratio in the spheroids assessed by Calcein AM (green, live cells) and ethidium homodimer dye (red, dead cells). (n = 6). Mean ± SEM, ** p < 0.01, One-way ANOVA followed by Tukey’s multiple comparison. **(G)** Representative western blots for cleaved caspase 3 in indicated AnS and AnR derivative cells after 24 hrs. in ultra-low attachment plates. Quantitation of cleaved caspase 3 normalized to β-actin shown below. **(H)** Flow cytometry analysis (left) and quantitation (right) of percent apoptosis determined using annexin V and PI staining in indicated AnS and AnR derivative cells after 24 hrs. in ultra-low attachment plates. (n = 3 biological replicates). Data are Mean ± SEM, *** p < 0.001, One-way ANOVA followed by Tukey’s multiple comparison. **(I)** Percent live cells in suspension of OV90 (left) and ID8 cells (right) derived from AnS or AnR (after 7 cycles of detachment-attachment) derivatives compared with the cells derived from ascites fluid at end point of mice that received OV90 and ID8 cells intraperitoneally. Cell viability in suspension was measured by trypan blue staining after 24 hours in suspension and plotted as percent survival in suspension (n = 2-5 replicates).

While the cell lines defined as AnR in Figure. 1A were resistant intrinsically, we asked whether cell lines defined as AnS could develop resistance to loss of attachment. To test this, we aimed to simulate a potential in vivo scenario of intraperitoneal and distant metastasis where cells are likely subjected to detachment stress followed by attached growth [38] [13]. Hence, we exposed the cells to repeated attachment-detachment conditions where cells were exposed to suspension cultures for 24 hrs and then allowed to expand in attached cultures (Figure 1B). We examined AnS cell lines that exhibited <100% viability at the first exposure to suspension (P1) (Figure 1B). 7/9 human cell lines and the mouse cell line tested, had significantly reduced cell death in suspension when subjected to sequential rounds of detachment followed by attachment growth (Figure 1C). Notably, the cell lines were able to maintain the acquired AnR phenotype through additional rounds of attachment-detachment (Figure 1C). For the remaining two cell lines tested (HEY, OVCAR3), while they appeared to become AnR (≥100% live cells) after 3-4 attachment – detachment cycles (Supplemental Figure 1C, HEY cells), they were unable to maintain this resistance and reverted to sensitivity through subsequent cycles of attachment-detachment.

We assessed if the AnR was due to changes in proliferation rate and/or population doubling times. Proliferative differences between the resistant, (AnR) or parental/sensitive, (anoikis sensitive: AnS,) cells were determined under both attached conditions and suspension culture conditions over a 10 day period. No significant differences in doubling time for the parental (AnS) and isogenic AnR cells in both 2D attached and suspension conditions were noted (Figure 1D). The parental AnS CAOV3 cells could not be assessed in suspension for 10 days due to extensive cell death. However, no significant differences were seen for the AnS and AnR pairs in attached growth over 10 days (Supplemental Figure 1D). Ki67 staining (Figure 1E) also revealed no significant differences between the AnS and AnR cells. In contrast, live/dead staining of both OV90 and CAOV3 parental AnS and isogenic AnR cells in suspension (Figure 1F), and analysis of cleaved caspase 3 from cells collected after 24 hrs in suspension revealed reduced cell death in the acquired AnR cells as compared to parental AnS cells (Figure 1G). These findings were further supported by flow cytometry analysis of annexin V and PI, which indicated a significant increase in the live/dead ratio (2.8 times higher for OV90 and 1.6 times higher for CAOV3) and reduced apoptosis (3.36 times lower in OV90 and 2.59 times lower in CAOV3) in AnR cells as compared to the parental AnS cells in suspension (Figure 1H).

To next evaluate if development of AnR as a result of cyclic exposure to suspension and attachment (Figure 1B) mimicked the development of AnR in vivo, we injected human OV90 cells and mouse ID8 cells i.p. into immunocompromised and immunocompetent mice respectively. Ascites fluid at the end point was used to collect cells that were then expanded in 2D for one passage followed by measurement of anchorage independent survival in suspension after 24 hrs. We find that both OV90 and ID8 cells from mouse ascites were resistant to cell death in suspension to similar extents as in vitro adapted AnR OV90 and AnR ID8 cells (Figure 1I). These data suggest that AnR cells adapted to anoikis in vitro show an AnR phenotype similar to in vivo adapted AnR cells from ascites.

### Acquired resistance to cell death in suspension is adaptive and reversible

To determine whether acquired AnR in vitro was due to clonal selection, genetic mutations, or non-genetic mechanisms, we used a modified Luria-Delbrück fluctuation analysis. Recent studies have adapted the classical Luria-Delbrück test to investigate cancer drug resistance by exposing single-cell derived clones to targeted therapy and analyzing fluctuations in surviving cell numbers to assess reversible switching between drug-sensitive and drug-tolerant states. [36–41] We asked if transient switching between cellular states could similarly drive AnR. We assessed survival fluctuations in suspension across single clones from the parental/AnS OV90 population (Figure 2A, n=60 single clones). Individual clones were expanded for 20 generations before determining live cell viability in suspension induced by detachment stress in polyhema-coated ULA plates (Figure 2A, B; P1 survival, grey bars). Clones were also maintained in 2D cultures, and viability of the expanded clonal populations was measured after three (P3) and six (P6) passages in 2D (Figure 2A, B). If clones switched between sensitivity (<100% survival) and resistance (≥ 100%), it would indicate non-heritable resistance. We observed no significant correlation in survival over time among clones (Figure 2C), which had doubling times ranging from 39 to 62 hrs (Supplemental Figure 2A, n=10 random clones), indicating the absence of fixed clonal states. Mean survival fractions (∼0.9) and interclonal fluctuations as quantified by the Coefficient of Variation (CV: 0.25-0.3) were consistent across different passages, (CV values for P1 = 0.283 ± 0.05, P3= 0.289± 0.05 and P6 = 0.259 ±0.05) with observed fluctuations exceeding those of the parental population (CV = 0.11 ± 0.04, obtained from n=11 biological repeats), where ± denotes the 95% confidence interval of the CV obtained from bootstrapping. If cells responded purely randomly to stress, fluctuations across clones would mirror population noise. However, the higher CV in clones suggests a memory effect or the presence of pre-stress states influencing detachment stress responses and anoikis resistance.

**Figure 2.**
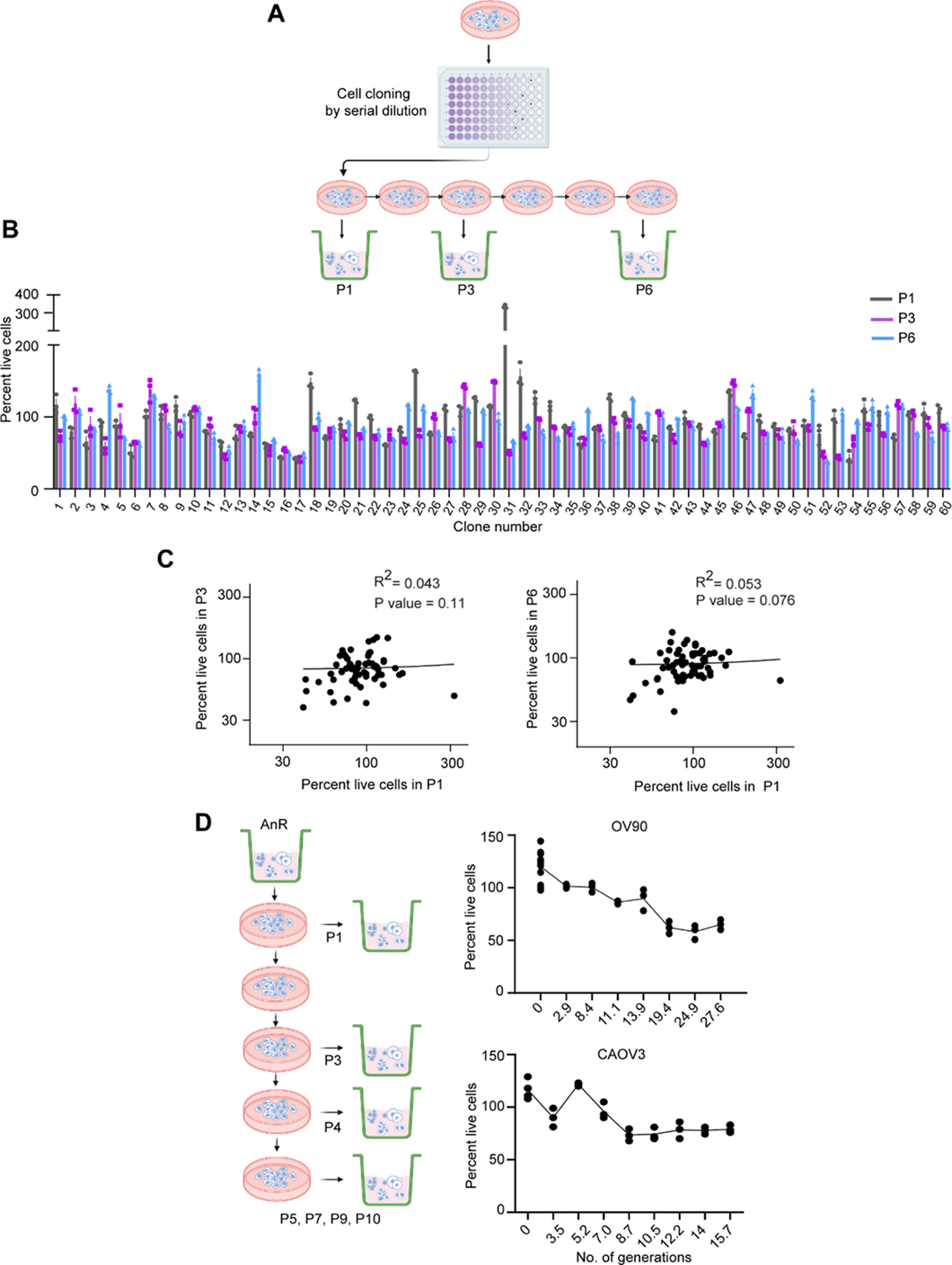
Acquired anoikis resistance represents a transient memory state in the population. **(A)** Scheme of single clone expansion from OV90 cells, followed by assessment of the survival of the individual clones upon suspension culture in ultra-low attachment plates for 24 hrs. in regular growth media at the indicated passages. **(B)** Percent survival of individual clones assessed according to the scheme in (A). (n = 60 clones assessed). **(C)** Linear regression correlation analysis of the survival of the individual clones in suspension P1 versus P3 (left) or in P1 versus P6 (right). **(D)** Experimental setup (left) to determine the stability of the adapted AnR cells maintained in attached conditions and tested for survival in suspension at the indicated passages after attached expansion. Percent survival in suspension plotted against the number of generations expanded in attached growth for OV90 and CAOV3 cells (right)

To test this memory effect, we evaluated the stability of the AnR state in OV90 and CAOV3 cells by propagating AnR cells in attached growth for several generations without suspension stress. When rechallenged with suspension stress, OV90 cells regained AnS after 11-14 generations and CAOV3 cells after 8-9 generations, approximating parental population sensitivity levels (Figure 2D). We applied previously developed analytical formulas [39–41] to predict what levels of fluctuations would be expected from switching between an anoikis-sensitive state (cell death in suspension) and an anoikis-resistant state (cell survival and proliferation in suspension). If *f* is the fraction of cells in the resistant state:

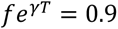

where 𝑇 = 24 ℎ𝑟 and growth rate 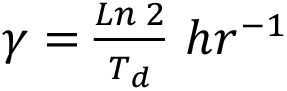 where 𝑇*_d_* is the cell doubling time of anoikis resistant cells in suspension that is experimentally determined. Thus, given a value of 𝑇*_d_*, the fraction of resistant cells can be computed from the above equation. Using a suspension doubling time of 100 hrs for OV90 in suspension (Figure 1D) results in f≈0.76. Model predicted fluctuations using equations derived previously, [39] individual clonal expansion in attached culture for 20 generations before the first survival test in suspension (Figure 2A,B P1), and a transient heritability of approximately 10-11 generations for the resistant state (Figure. 2D), were <0.01 and 30-fold lower than observed values (0.25-0.3) for biologically relevant T_d_ (≥38 hrs., OV90 2D doubling time, Figure.1D). Thus, reversible switching between these two phenotypic states cannot explain the observed clone-to-clone variations in surviving cells.

To uncover mechanisms underlying interclonal fluctuations, we calculated the effective growth rate during the first suspension test using 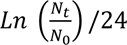, where N_0_ and N_t_ are cell numbers at the start and end of the 24-hr suspension period. Clones exhibited significant variation in growth rates, averaging -0.004 hr ^-1^ and a CV of 300%. Approximately two-thirds of clones showed negative growth rates (N_t_<N_0_), while one-third had positive growth rates (N_t_ > N_0_), with four clones doubling in suspension (doubling time: 45 hrs.) similar to measurements in 2D cultures. These high clonal growth rate variations suspension stress could not be captured by a simple two state model. Together with the lack of concordance between the predicted and observed clonal fluctuation, these data suggest a continuum of cellular states but are however consistent with non-genetic effects on adaptation of cells as they develop resistance to the biological stress of loss of attachment (AnR). Non-genetic adaptation was confirmed by whole exome sequencing. The total number of genes that were mutated in any given replicate was found to be 2860. However, no significant differentially mutated genes (using a p value of p < 0.05 and Fisher’s exact test) were found between the two isogenic populations of cells (Top 50 mutated genes based on total number of mutations present in OV90 cells is shown in Supplemental Figure 2B). Thus, acquired resistance was not due to clonal selection of an anoikis-resistant subpopulation that preexisted, but rather represents the development of an anoikis tolerant state.

### Adapted anoikis tolerant ovarian cancer cells are more chemoresistant and metastatic in vivo

Prior studies have implicated anoikis resistance as a phenotype of metastatic and chemo resistant cells. [5, 42–44] Given that reversion to the original AnS state occurred in the absence of stress (Figure 2D) we tested if such an acquired ‘tolerance’, rather than permanent/ (intrinsic) resistance, was sufficient to alter in vitro properties associated with tumorigenicity. A transwell migration assay revealed that OV90-AnR cells (derived from P7) were able to undergo significantly higher migration through fibronectin (Figure 3A) as compared to the parental population (P0), and as compared to cells expanded after a single exposure to suspension culture (P1, Figure 3A). Increased migration of AnR cells was seen not just in OV90 cells, but also in CAOV3 cells (Figure 3A). The increased migration of adapted AnR cells was seen across additional isogenic pairs tested including EOC15, p151 and OVCA420 (Supplemental Figure 3A). All tested pairs revealed increased migration for the AnR derivatives only, demonstrating increased motility as a feature of acquired AnR. Assessing the chemosensitivity of the cells to standard of care chemotherapeutics including cisplatin, paclitaxel, and doxorubicin revealed that AnR cells across cell line models had significantly higher IC50 concentrations for paclitaxel (Figure 3B). Increased IC50 for paclitaxel was also seen for the AnR cells in suspension cultures (Supplemental Figure 3B). However no significant differences were observed for doxorubicin and cisplatin (Supplemental Figure 3C) in an acute response test.

**Figure 3.**
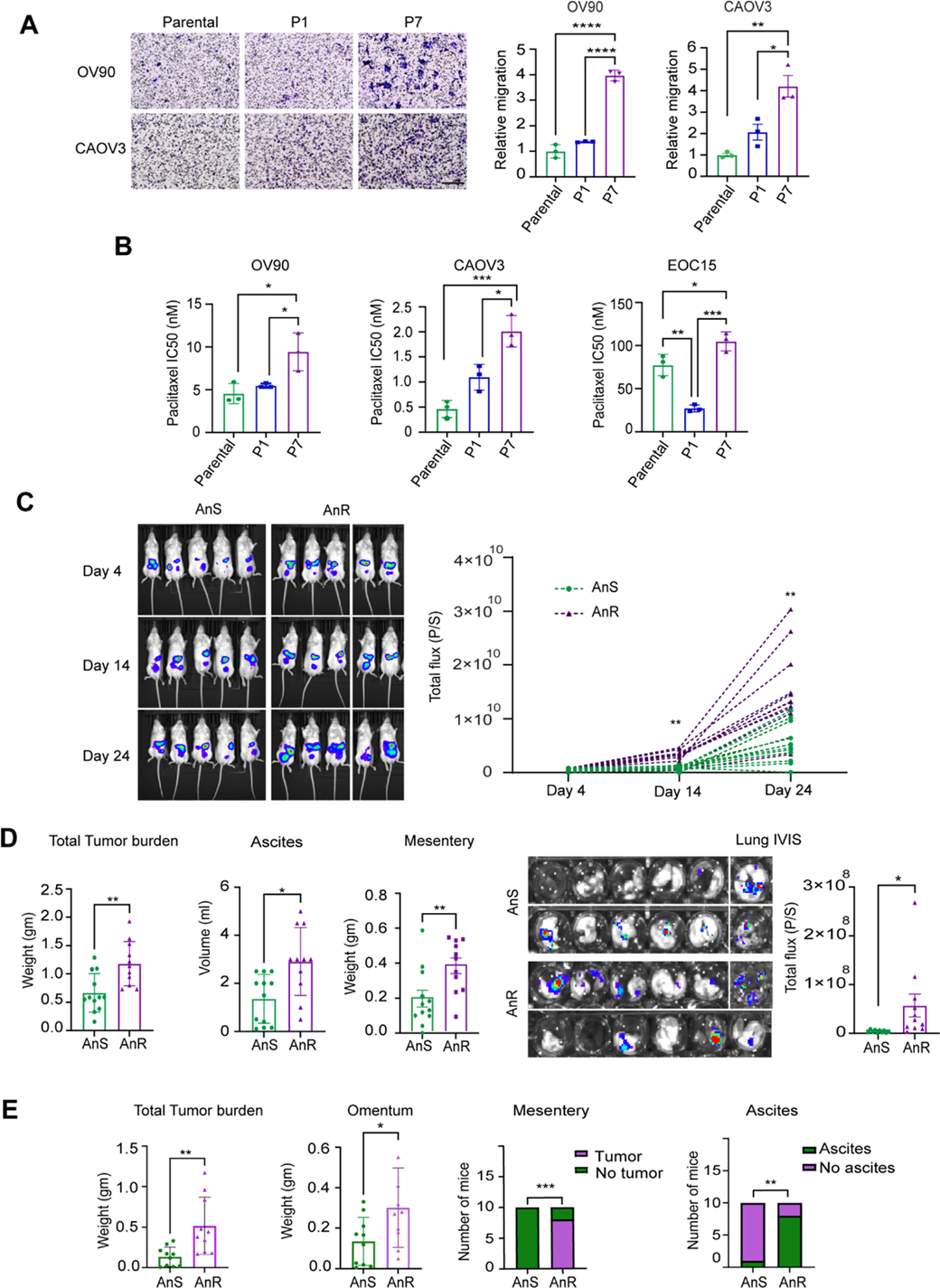
Acquired anoikis resistance leads to increased in vitro migration, chemoresistance and in vivo intraperitoneal growth and metastasis. **(A)**. Representative images (left) and quantitation (right graph) of indicated parental, P1 (cells expanded after one exposure to suspension culture) and P7 (anoikis resistant/AnR) cells on fibronectin-coated transwell filters after migration for 24 hrs. (n = 3). Data are mean ± SEM. ns p > 0.05, * p < 0.05, ** p < 0.01, **** p <0.0001. One-way ANOVA followed by Tukey’s multiple comparison. **(B)**. IC50 to paclitaxel of OV90, CAOV3 and immortalized EOC15 cells determined under steady state attached conditions of indicated cells after 72 hrs. using an SRB assay. Data are mean ± SEM. ns p > 0.05, * p < 0.05, ** p < 0.01, **** p <0.0001. One-way ANOVA followed by Tukey’s multiple comparison. **(C)**. Whole body luminescence (BLI) (left) representative images and quantification (right) of total flux over time of NOD-SCID mice injected with 5 million live OV90 parental (P0/AnS) or anoikis resistant (P7/AnR) cells. Data are mean ± SEM. ns p > 0.05, ** p < 0.01, *** p <0.001, Two-way ANOVA followed by Tukey’s multiple comparison. **(D).** Total tumor weight (gms), total ascites volume (ml) and mesenteric weight from mice receiving P0 (AnS) or P7 (AnR) OV90 cells as indicated, analyzed between days 39-40. Luminescence images of explanted whole lungs (left) and quantitation of total luminescence flux of the lungs (right) from mice receiving either P0 or P7 OV90 cells at days 39-40. n =12 for AnS/P0 and 11 for AnR/P7. All data are Mean ± SEM; * p < 0.05, ** p < 0.01, unpaired t test. **(E)**. Analysis of total tumor weight, weight of the omentum, no. of mice with tumors in the mesentery and no. of mice with retrievable ascites from C57BL6 mice, i.p. injected with either parental AnS (P0) murine ID8 or AnR (P8) ID8 cells between day 80-82. n =10. All data are Mean ± SEM; * p < 0.05,** p < 0.01, ***p<0.001, unpaired t test.

In order to assess if acquired AnR was sufficient to increase intraperitoneal (i.p.) growth and metastasis of tumor cells, we injected OV90 luciferase expressing parental cells (parental/P0), or cells expanded under attached conditions after development of AnR (AnR/P7), into the peritoneal cavity of NOD-SCID mice. Whole body bioluminescence imaging (BLI) revealed a significant increase over time in i.p. tumor burden in mice receiving AnR cells as compared to mice receiving parental (P0) cells (Figure 3C). Since ascites accumulation, which began at ∼day 25, precludes reliable BLI, BLI data are presented only until day 24 (Figure 3C). The study was terminated at day 39-40 when mice from the P7/AnR group were found to be moribund. End point total tumor weight in the peritoneal cavity of mice receiving AnR cells was twice as much as compared to the mice receiving the parental population (AnS) (Figure 3D), concomitantly with higher volume of ascites, and tumor burden in the mesentery (Figure 3D). Strikingly, animals receiving AnR cells (P7) showed higher metastatic growth in the lungs as determined by BLI imaging of explanted lung tissues as compared to the parental cells (P0) (Figure 3D).

Since the in vitro derived AnR cells mirror suspension survival of in vivo derived AnR cells of human (OV90) and mouse (ID8) origin (Figure 1I), we tested if acquired AnR was sufficient to increase i.p. growth and metastasis of tumor cells even in the presence of the immune system. We injected ID8 parental cells (parental/P0) or in vitro adapted AnR cells (AnR/P7) into the peritoneal cavity of C57BL6 mice. The study was terminated when mice from the P7/AnR ID8 group exhibited signs of being moribund. Adapted ID8 AnR cells produced significantly higher disease burden as evident from the higher overall measurable tumor burden (Figure 3E) including in the omentum and mesenteric regions and from the number of mice with measurable ascites (Figure 3E). Lung metastasis, however, was not evident from pathological assessments in the ID8 model. These data demonstrate that repeated cycles of exposure to suspension stress followed by attached growth leads to adaptation that is sufficient to promote aggressive disease in vivo mimicking disease spread seen in advanced stage OC patients.

### Development of adaptive anoikis resistance is concomitant with transcriptional reprogramming over time

We next assessed the transcriptional changes in response to suspension stress and during recovery periods of attached growth. Bulk RNA sequencing of OV90 and CAOV3 cells undergoing adaptation at various suspension and recovery time points revealed that the number of Differentially Expressed Genes (DEGs, defined by padj <= 0.05, L2FC >1.5) after the first exposure to suspension stress (P0 versus P1 after 24 hrs. in suspension) was 1,011 for OV90 and 362 for CAOV3, of which a total of 148 were upregulated and 863 downregulated for OV90 and 171 and 191 for CAOV3 respectively (Figure 4A-C, Supplemental Figure 4A-C). In the case of OV90, 74 of the 148 upregulated genes were upregulated only during the first exposure to stress (P0-P1) (Figure 4A). As cells adapted to stress in subsequent cycles, the total number of DEGs decreased over time, with more genes downregulated than upregulated in both OV90 and CAOV3 cell lines (Figure 4A-B, Supplemental Figure 4A-B). The highest number of unique upregulated genes was observed from P0 to P7 (208 genes, OV90), and during the transition to resistance in OV90 cells (P0 and P3 time points; 165 genes). Most of the unique downregulated genes were found between P0 vs P7 (suspension AnR) and P0 vs P6 (attached AnR), with 375 and 422 genes, respectively. 225 of these genes were consistently downregulated over time (Figure 4B last column). For CAOV3, the comparison between P0 and P6 (2D comparisons of AnS and AnR), had a total of 131 upregulated genes and 213 downregulated genes. 29 upregulated and 115 downregulated were unique to this comparison. (Supplemental Figure 4B-C). The P0 vs P7 comparison, gave the greatest number of uniquely up- and downregulated genes (110 and 379, respectively) in CAOV3 (Supplemental Figure 4A-C), similarly to OV90. Thus, while the number of DEGs per time point varied between the cell lines (Figure 4A-C, Supplemental Figure 4A-C) the number of differentially downregulated genes exceeded the number of differentially upregulated genes in both models. The magnitude and timing of changes also differed between cell lines suggesting cell line-specific adaptation kinetics. Together these data demonstrate significant transcriptional changes occurring after the first exposure to stress, with only a subset of the same changes maintained in later stages of adaptation.

**Figure 4.**
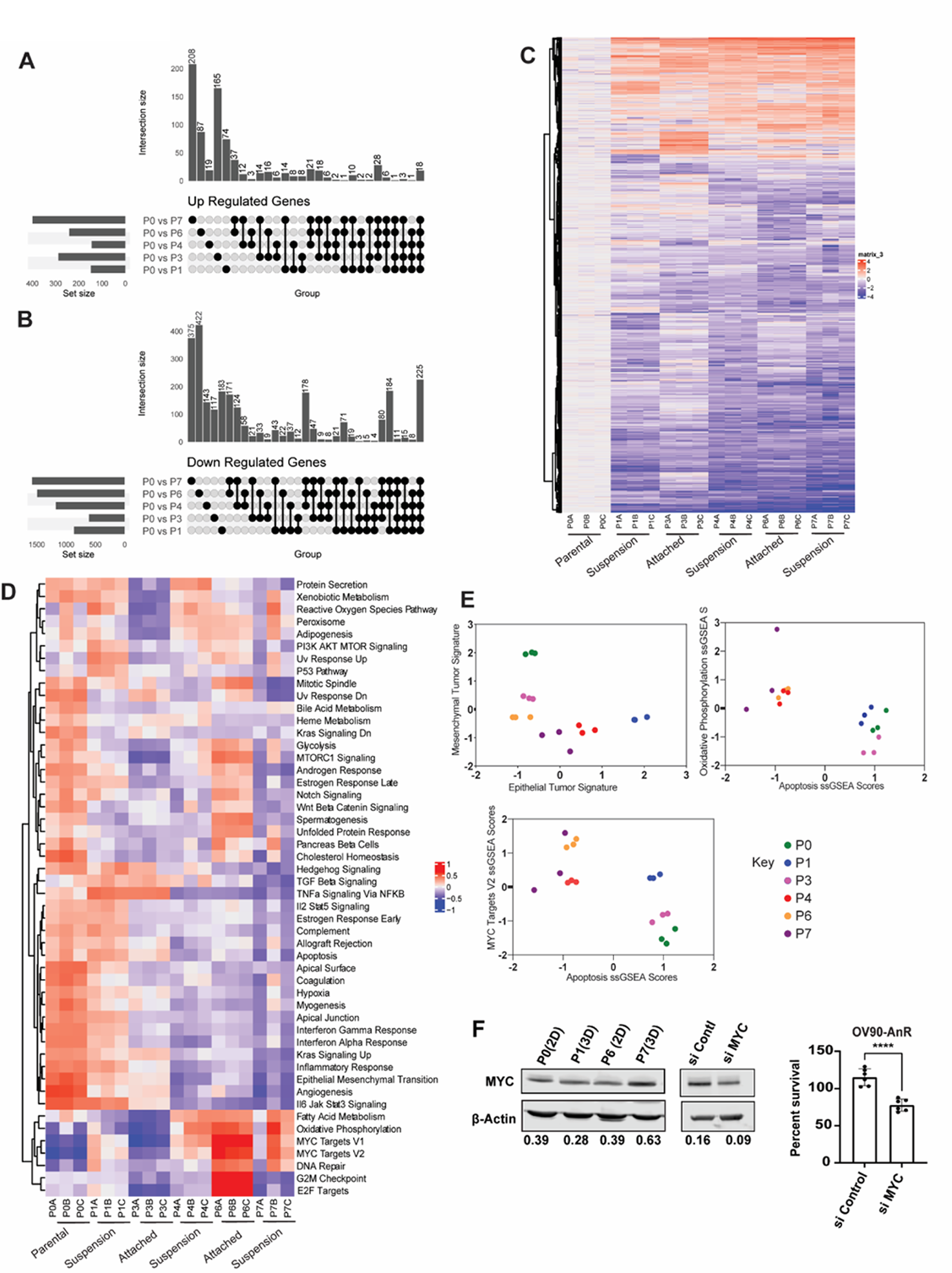
Transcriptional changes and pathway alterations during adaptation to anoikis over time. **(A)**. UpSet plots highlighting the number of Up Regulated Genes (A) (p value < 0.05 and L2FC >1.5) and Down Regulated genes **(B)** (p value < 0.05 and L2FC < -1.5) in OV90 across the time point comparisons P0 vs P1, P0 vs P3, P0 vs P4, P0 vs P6, and P0 vs P7 as indicated. **(C)** Heatmap for the individual samples using Log2FC as calculated for individual biological replicates for parental (P0 cells) (P0A-C), P1 (P1A-C), P3 (P3A-C), P4 (P4A-C), P6 (P6A-C), and P7 (P7A-C). Respective attached and suspension time points are indicated. Heatmap generated by clustering analysis of DEGs across samples. The Log2 fold changes of each gene were clustered based on euclidean distance. **(D)**. Heatmaps for the individual OV90 samples using GSVA normalized enrichment scores (NES) for passage 0 (P0A-C), 1 (P1A-C), 3 (P3A-C), 4 (P4A-C), 6 (P6A-C), and 7 (P7A-C). The hallmarks were clustered based on euclidean distance. **(E)** Scatterplot showing the NES for samples on a 2-dimensional epithelial mesenchymal plane (left) , Scatterplot showing the NES for apoptosis and the NES for Oxidative Phosphorylation (right) across time points for each sample in the replicates for OV90, and Scatterplot showing the NES for apoptosis and the NES for MYC Targets V2 (bottom) for each time point for OV90. **(F)** Representative western blot (left) for c-MYC in indicated OV90 cells. Quantitation of c-MYC normalized to β-actin shown below. (n = 2). Representative western blot (middle) for c-MYC in P7/AnR OV90 cells following transient knockdown using siRNA against c-MYC. Quantitation of c-MYC normalized to β-actin shown below. Percent live cell count (right) of OV90 from P7 after 24 hours in suspension assessed by Trypan blue exclusion assay following transient knockdown of c-MYC. Data are Mean ± SEM; ****p < 0.0001., unpaired t test.

Gene Set Variant Analysis (GSVA) of the different time points using the Human MSigDB Hallmark Collections revealed a subset of hallmarks in OV90 cells that were positively enriched during the adaptation cycles, such as those associated with ‘TGF beta signaling’, ‘Hedgehog signaling’ and ‘TNFa signaling via NFkb’ (Figure 4D). Later passages showed reduced apoptosis (Figure 4D, E) and the cells alternated between epithelial and mesenchymal states before converging toward a hybrid EM state once adapted (Figure 4E). Additional pathways were more specific to attached growth conditions in the adapted resistant stages in OV90 cells, such as mTORC1, Notch, and Wnt beta catenin (Figure 4D, P6). GSVA analysis in CAOV3, which is more anoikis sensitive than OV90 (Figure 1), has a different mutational background that OV90 (depmap.org) and reverted back to sensitivity sonner than adapted OV90 ARE cells (Figure 2D), identified overlapping and some distinct enrichments as compared to OV90. Hallmarks including ‘TGF-β signaling’, ‘Hypoxia’ and ‘Hedgehog signaling’ and “TNFa signaling via NFkb’ that were enriched during adaptation in OV90 cells, remained enriched in CAOV3 even at later time points (Supplemental Figure 4D) suggesting differences in the timing and maintenance of the pathway activation between the two models. Both cell lines however showed common enrichments in ’Oxidative Phosphorylation’ and ’MYC Targets’ in adapted cells (Figure 4E, Supplemental Figure 4D). Increase in *MYC* RNA levels (Supplemental Figure 4E) was also manifested at the protein level (Figure 4F), with lowering MYC in the adapted AnR OV90 cells using siRNA leading to reduced survival in suspension (Figure 4F, right). Together, these data highlight common mechanisms enriched during, and in adapted cells, with additional pathways altered at different times between models during adaptation.

### Adapted anoikis resistant cells depend on oxidative phosphorylation for survival

Changes in the ‘Oxidative Phosphorylation’ pathway in adapted anoikis resistant cells was a hallmark in both OV90 and CAOV3 (Figure. 4D,E and Supplemental Figure 4D). We thus tested for direct effects on mitochondrial respiration in anoikis adapted cells. We first measured parameters of mitochondrial function by assessing respiration using extracellular flux analysis with a mitochondrial stress test (seahorse XF96 assay) in both OV90 and CAOV3 cells (Figure 5A). When comparing parental cells (P0) to cells after exposure to one round of suspension stress (P1), or to adapted AnR/P7 cells expanded in attached conditions, we find that AnR cells from both OV90 and CAOV3 cells exhibited higher oxygen consumption and extracellular acidification rates (OCR and ECAR respectively) at baseline compared to parental cells (P0) and cells from P1 (Supplemental Figure 5). ATP-dependent OCR, measured by adding oligomycin to inhibit Complex V (ATP synthase) of the electron transport chain (ETC) to the culture (Figure 5A), revealed significantly higher ATP-dependent OCR in P7/AnR cells (mean=121.5 pmol/min/cell -/+) compared to parental P0 cells (84.8 pmol/min/cell -/+) and P1 (83.4 pmol/min/cell -/+) in OV90 cells (Figure 5A). In CAOV3 cells, ATP-dependent OCR in P7 was also higher compared to P0 and P1 (Mean ATP production in Parental, P1 and P7 was 61.8, 59.2, and 98.9 pmol/min/cell respectively) (Figure 5A). Spare respiratory capacity (SRC) is a parameter representative of the mitochondrial ability to produce energy through respiration beyond the amount needed for basal cellular maintenance and has been shown to be increased during malignant transformation and tumor invasion. [45] SRC determined by adding FCCP (an uncoupler of mitochondrial oxidative phosphorylation) that shortcuts the ETC (Figure 5A) and increases OCR to its maximal level revealed that SRC was significantly higher in AnR/P7 cells compared to parental (P0) and P1 cells in both OV90 and CAOV3 models (Figure 5A).

**Figure 5.**
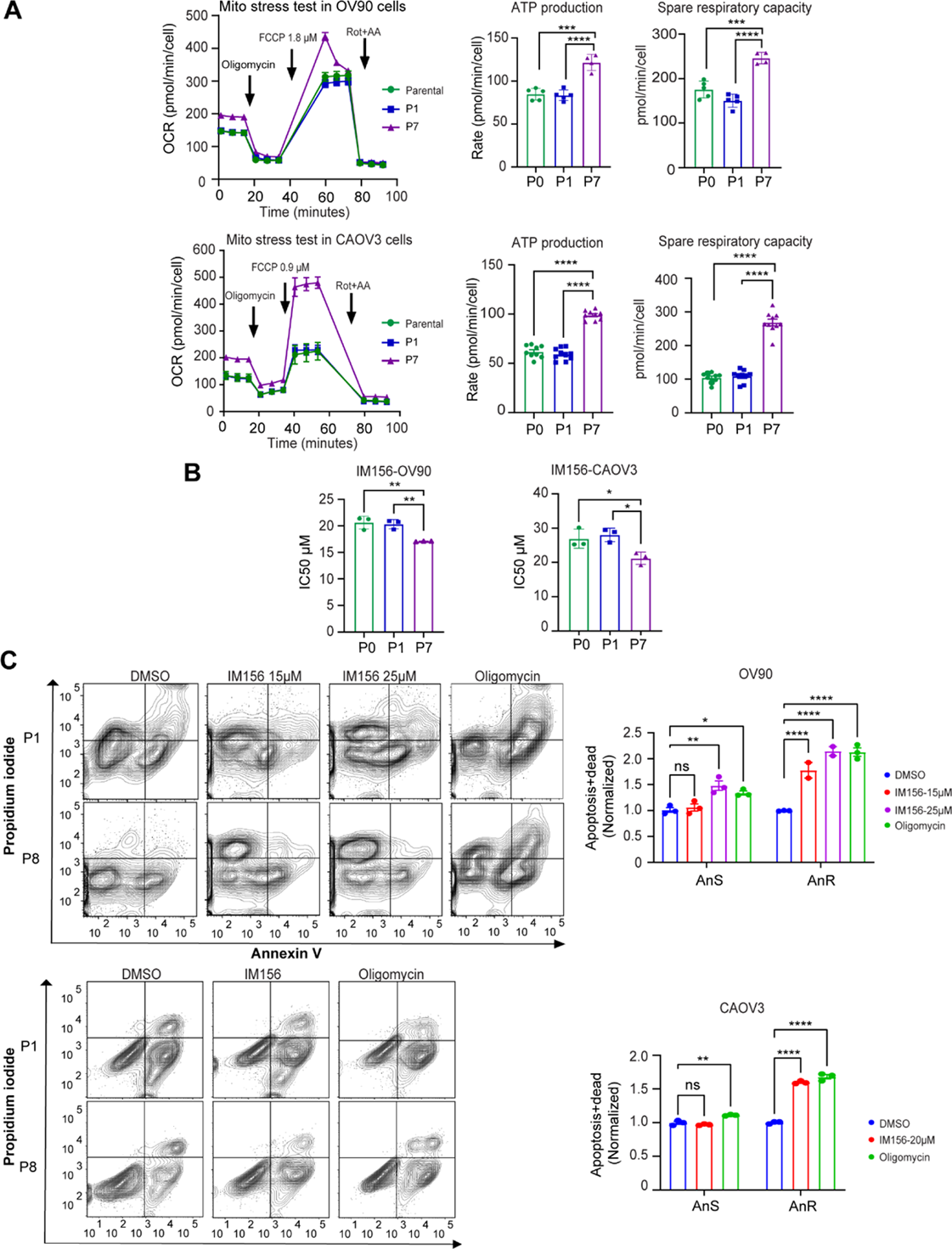
Adapted anoikis resistant cells are dependent on oxidative phosphorylation for enhanced survival upon loss of attachment. **(A)**. Representative Oxygen consumption rate (OCR) (left) of cells from either parental AnS (P0), cells exposed to one cycle of suspension culture (P1) or adapted AnR (P7) cells measured using Seahorse XF96 mito stress test under attached conditions at baseline or after addition of Oligomycin (1.5 µm), FCCP (1.8 µm for OV90 and 0.9 µm for CAOV3), and Rotenone-Antimycin A (0.5µm), ATP production (middle) and spare respiratory capacity (right) calculated from OCR and ATP production for indicated OV90 and CAOV3 cells (n = 2). Data are normalized using SRB and Data are mean ± SEM. ns p > 0.05, *** p < 0.001, **** p <0.0001. One-way ANOVA followed by Tukey’s multiple comparison. **(B)** IC50 of OV90 or CAOV3 cells to IM156 determined under attached conditions after 96 hrs. using an SRB assay for cells from the indicated passages (n =3). Data are Mean ± SEM. ns p > 0.05, * p < 0.05, ** p < 0.01, One-way ANOVA followed by Tukey’s multiple comparison. **(C)** Flow cytometry analysis and quantitation (adjacent panel) of the apoptotic and dead cell population determined using Annexin V and propidium iodide staining of cells from indicated passages of either OV90 (top) or CAOV3 (bottom) treated for 72 hours with DMSO, IM156, and Oligomycin in ultra-low attachment plates. Data are normalized to the respective DMSO controls (n = 3). Data are Mean ± SEM. ns p > 0.05, * p < 0.05, ** p < 0.01,**** p <0.0001, two-way ANOVA followed by Tukey’s multiple comparison.

Since adapted anoikis resistant cells showed higher SRC, we compared the sensitivity of parental (P0) and P1 cells and AnR /P7 cells to the biguanide IM156, an AMPK activator and Complex I inhibitor currently in clinical trials. [46] Remarkably, we found that adapted AnR cells from P7 for both OV90 and CAOV3 cells showed significantly lower IC50 to IM156 compared to parental and P1 cells. (OV90 mean IC50; parental= 20.6, P1= 20.3, P7=17.1µM and CAOV3 mean IC50; parental= 26.9, P1= 28, P7= 21.2 µM, Figure 5B). Moreover, a sub IC50 dose of IM156 induced cell death in suspension (anoikis) to a significantly greater extent in the adapted AnR/P7 cells as compared to the parental counterpart (Figure 5C). Similarly, inhibition of the ETC complex V using oligomycin was able to re-sensitize and induce anoikis to a significantly higher degree in the adapted AnR P7 cells as compared to parental cells (Figure 5C). These data demonstrate that adapted AnR cells have developed an enhanced mitochondrial capacity to support their superior survival in suspension and indicate that blocking OXPHOS in these cells could reverse such an adaptive phenotype.

### Adapted anoikis resistant cells evade immune surveillance

Bulk RNA seq results also revealed perturbation in pathways that may impact immune recognition of the adapted AnR cells compared with their sensitive counterpart. Of note, the inflammatory response pathway was enriched early during adaptation (P0 (parental), P1 and P3 (collected after 1 and 3 cycles of detachment respectively) in comparison to later stages of adaptation P4, P6, and P7 (collected after 4,6, and 7 cycles of detachment respectively) in OV90 cells (Figure 6A). We also examined expression of HLA genes, as downregulation of MHC-I antigen presentation is a known mechanism of immune evasion in multiple cancers [47] and has been associated with anoikis resistance in other cancers. [44] We found that HLA-B as part of major histocompatibility complex I (MHC-I) but not HLA-A and C was downregulated over time during development of adaptive AnR in the attachment-detachment cycles in OV90 cells (Figure 6B). Protein levels of MHC-I protein were also lower in adapted P6 and P7 cells compared with P0 and P1 cells (Figure 6C). Additionally, TAP1 (Transporter associated with antigen processing 1) that is involved in MHC-I antigen presentation, was absent in adapted AnR cells (Figure 6C). TAP1 and MHC-I play crucial roles in tumor antigen recognition by cytotoxic CD8+ T cells. [48] We thus hypothesized that adapted AnR cells that have reduced expression of MHC-I and TAP1 could be less susceptible to immune mediated killing compared to their AnS counterparts. We isolated and activated CD8+ T cells from PBMCs derived from healthy individuals (Figure 6D) and added them to OV90 and CAOV3 cells from P0, P1, and P7 cells cultured in regular attached conditions (Figure 6D). We found a time dependent decrease in cell viability of the P0 OV90 (parental population) as compared to the adapted AnR OV90 cells (Figure 6D). For CAOV3 cells we observed a much stronger and faster T cell killing response compared to OV90s, as at 24 hours, only 10% of viable cells remained in the parental AnS cells (P0, P1). In contrast, 40% of adapted AnR cells (P7) survived (Figure 6D). To measure apoptosis of tumor cells induced by CD8+ T cells, we examined cleaved caspase 3 (CC3) after 48 hours of co culture of OV90 cells with T cells and 12 hrs. of co-culture for CAOV3 cells due to their higher sensitivity (Figure 6E). We find significantly higher CC3 in AnS cells (P0, P1) as compared to the isogenic AnR cells (P7) (Figure 6E). These data suggest that adaptation to detachment-induced cell death in ovarian cancer cells facilitates immune evasion concomitant with reduced antigen presentation likely through the observed downregulation of MHC-I and TAP1.

**Figure 6.**
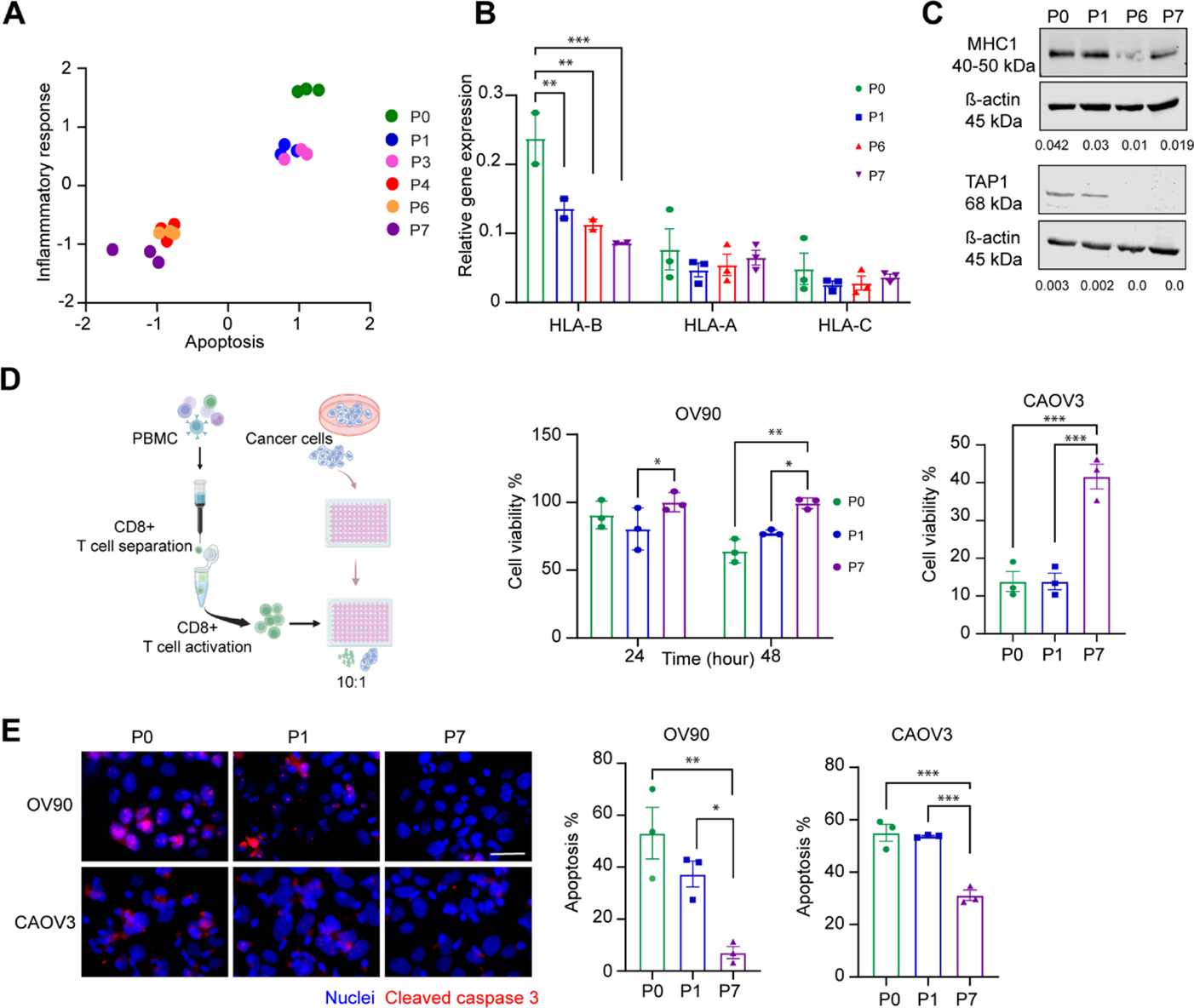
Adapted anoikis resistant cancer cells evade killing by cytotoxic T cells. **(A)** Normalized Enrichment score (NES) for Hallmark inflammatory response pathway plotted against Hallmark apoptosis in OV90 cells derived from P0 (parental), P1, P3, P4, P6, and P7 (after 1 to 7 cycles of suspension culture for 24 hours respectively) n =3 replicates. **(B)** Semi qRT-PCR analysis of HLA-A, B, and C mRNA expression in indicated anoikis sensitive (P0, P1: after one round of 24 hr suspension stress) and isogenic anoikis resistant (P6, P7) OV90 cells. n =2-3 replicates. Data are mean ± SEM. ns p > 0.05 ** p < 0.01, *** p < 0.001. Two-way ANOVA followed by Tukey’s multiple comparison. **(C)** Representative western blot for TAP1 (lower) and MHC-I (upper) in indicated AnS and AnR derivative cells. Quantitation of TAP1 and MHC-I normalized to β-actin shown below. (n = 2). **(D)** Schematic (left) of immune mediated tumor killing assay. CD8+ T cells were isolated from PBMCs and activated for 48 hours then co cultured with either OV90 or CAOV3 cells in the attached condition. Percent cell viability (right) of indicated OV90 and CAOV3 cells measured by trypan blue staining after incubation with activated CD8+ T cells (24 and 48 hours for OV90 and 24 hours for CAOV3). n =3, Data are mean ± SEM. ns p > 0.05, * p < 0.05, ** p < 0.01, *** p < 0.001. Two-way ANOVA for OV90 and One-way ANOVA for CAOV3 followed by Tukey’s multiple comparison. **(E)** Representative widefield immunofluorescence images (left) of Cleaved Caspase 3 (red) and DAPI (blue), and quantitation of percent apoptosis (right) from indicated OV90 and CAOV3 cells co-cultured with activated CD8+ T cells (48 hours for OV90 and 12 hours for CAOV3 cells). Data are mean ± SEM. ns p > 0.05, * p < 0.05, ** p < 0.01, *** p < 0.001, **** p <0.0001, One-way ANOVA followed by Tukey’s multiple comparison. Scale bar: 50 µm. (n = 3 biological trials, each trial includes quantitation of a minimum of four 60X fields).

### Inhibiting the transcriptional Mediator kinase CDK8/19 suppresses anoikis resistance in vitro and intraperitoneal growth in vivo

Transcriptional changes were both overlapping and different between the two cell line models including changes in the timing of the pathways being enriched. We thus tested the effects of inhibiting the CDK8/19 Mediator kinase, a pleiotropic regulator of transcriptional reprogramming, on development of adaptive AnR. For this, we used SNX631 as a selective inhibitor of CDK8/19. [49] We first assessed sensitivity of a panel of intrinsically AnS cell lines (OV90, CAOV3, EOC15, and OVCA420) to SNX631 over a 7 day treatment course, as CDK8/19 inhibition has been reported to be cytostatic in a subset of cancers. We find the IC50s of the cell lines to range between 2.16 µM for OV90 to 4.1 µM for OVCA420 (Figure 7A), which is over 2 orders of magnitude higher than the IC50 of SNX631 in a cell-based assay for CDK8/19 inhibition (11 nM). [49] Hence, CDK8/19 activity did not appear to be required for proliferation and steady state growth of the tested cell lines in vitro. Given the stress associated with matrix detachment, we tested the effect of SNX631 on adaptation to anoikis resistance. For this, we chose a 500 nM concentration of SNX631 which does not inhibit cell growth but is sufficient for complete CDK8/19 inhibition in a cell-based assay [50] and reducing the phosphorylation of STAT1 at S727 (Figure 7A ), which is phosphorylated directly (but not exclusively) by CDK8/19. [51, 52] We found that treatment of OV90, OVCA420 and immortalized EOC15 cells with 500 nM SNX631 during cyclic attachment-detachment culture cycles prevented the development of AnR, with the mean percent live cells reaching a maximum of only 53.28 %, 63.7 %, or 67% respectively for each of the models. while DMSO-treated control cells reached or exceeded 100% (Figure 7B). These data demonstrate that SNX631 prevented the development of AnR across all OC models tested. To assess whether global downregulation of the overall transcription rather than transcriptional reprogramming is sufficient to block the development of anoikis resistance we treated OV90 cells during cyclic culture of attachment -detachment with THZ1 which is a covalent CDK7 inhibitor that also targets CDK12 and was shown to globally downregulate transcription. [53] Prior studies established that THZ1 potently inhibits CDK7 with an IC50 of ∼3.2 nM in vitro, with >50% inhibition in cellular assays achieved in the range of 10–40 nM. [53] We find that in contrast to CDK8/19 inhibition, a sublethal concentration of THZ1 that is known to be effective at CDK7 inhibition [53] and chosen based on IC50 in OV90s (Supplemental Figure 6A) had no impact on development of AnR (Supplemental Figure 6B). These data suggest selective dependence on CDK8/19 rather than overall transcription during development of AnR.

**Figure 7.**
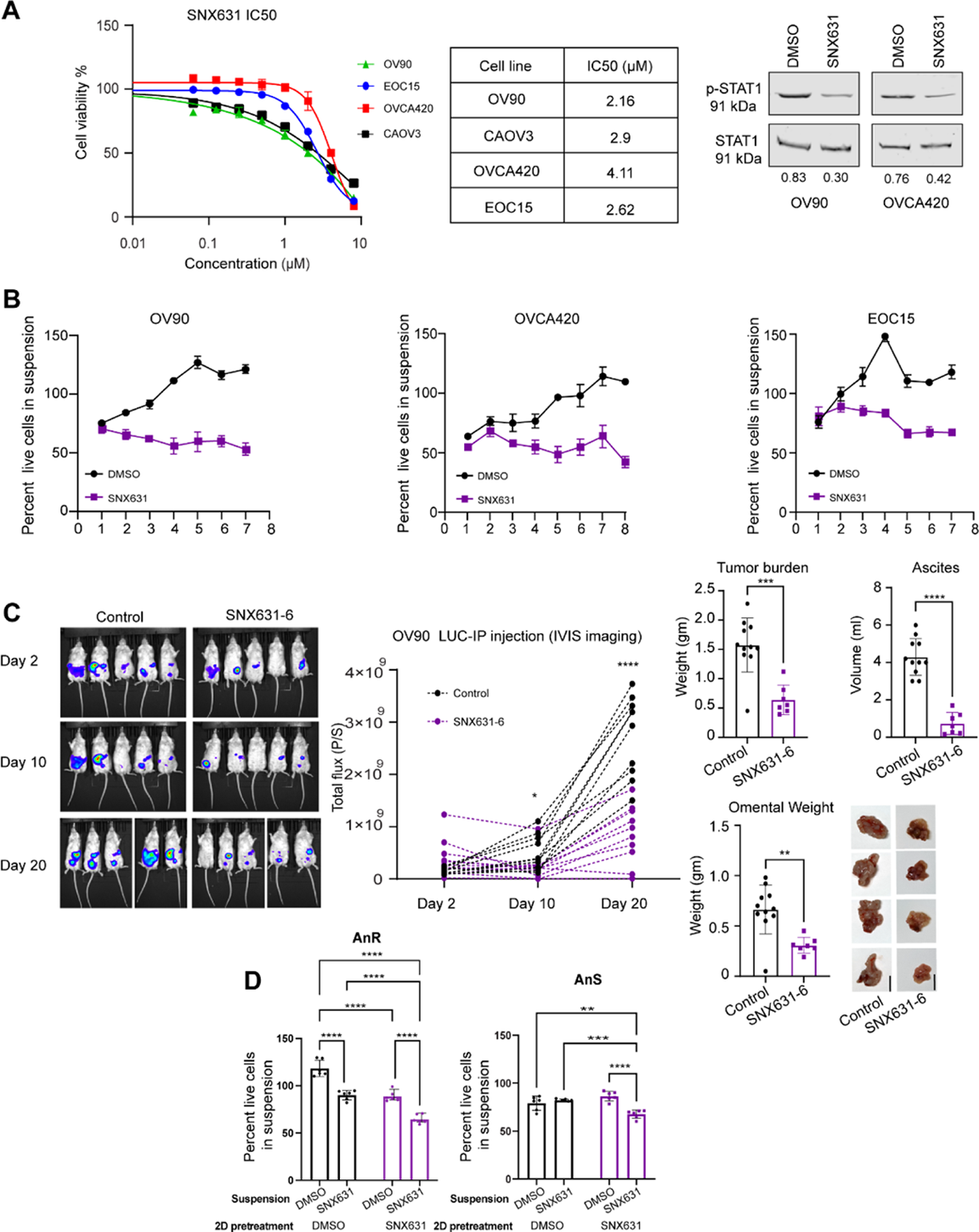
CDK8/19 Mediator kinase inhibition prevents development of anoikis resistance and i.p. xenograft tumor growth. **(A)** IC50 of indicated cell lines assessed after 7 days of incubation with either vehicle or SNX631 by SRB assay which was used to calculate IC50 (middle). pSTAT1 and total STAT1 (right) in OV90 and OVCA420 cells in response to 500nM SNX631 treatment for 24 hrs. **(B)** Percent survival of indicated anoikis sensitive cell after 24 hours in suspension following cycles of gain and loss of attachment as in Fig 1B, either with vehicle (DMSO) or in the presence of 500 nM CDK8/19 Mediator kinase inhibitor (SNX631) (n = 3). **(C)** Whole body luminescence (BLI) representative images (left) and quantification (middle) of total flux over time of NSG mice injected with 5 million live OV90 (P0) cells receiving either control diet or SNX631-6 medicated diet for the duration of the study. (n =11-12). Data are Mean ± SEM. ns p > 0.05, * p < 0.05, **** p <0.0001, two-way ANOVA followed by Tukey’s multiple comparison. Total tumor weight, ascites volume, and omental weight, with example pictures (right), from mice receiving either a control diet or SNX631-6 medicated diet analyzed at the end point on day 41. (n =7-11). Data are Mean ± SEM. ** p < 0.01, *** p < 0.001, **** p <0.0001 Student t-test. **(D)** Percent survival of either AnR OV90 or parental AnS OV90 pretreated in 2D cultures for 96 hours with either vehicle (DMSO) as control or 500 nM SNX631 followed by 24 hours in suspension, either in the presence of vehicle (DMSO) or 500 nM SNX631 (n =6/condition). Data are Mean ± SEM. ns p > 0.05, * p < 0.05, **** p <0.0001, two-way ANOVA followed by Tukey’s multiple comparison.

Since development of AnR in vitro, mimicked development of AnR in vivo (Figure 1I) to test whether CDK8/19 inhibition also affects in vivo intraperitoneal (i.p.) growth of OC cells, we injected OV90 parental/AnS luciferase expressing cells into the peritoneal cavity of NSG mice. Mice were randomized in two groups, one receiving SNX631-6 (an equipotent analog of SNX631 and a clinical drug candidate [54]), administered in medicated diet (at 350 ppm [54]); the second group received control diet. Whole body bioluminescence imaging (BLI) revealed a significant decrease in overall i.p. tumor burden over time in mice receiving SNX631-6 (Figure 7C). The study was conducted until day 41, with BLI data shown until day 20 (Figure 7C) as ascites buildup in the control group starting at ∼week 4 post tumor cell injection interfered with accurate luciferase detection in animals. Endpoint total tumor burden, ascites volume and omental tumors were significantly reduced in size, volume and weight in mice receiving SNX631-6 (Figure 7C). These results indicate that inhibition of CDK8/19-regulated transcriptional reprogramming and prevented and suppressed AnR in vitro and intraperitoneal tumor growth and metastasis in vivo.

We next wanted to test if SNX631 could also resensitize AnR cells to anoikis. Since OC cells tested were not sensitive to SNX631 in steady state 2D growth (Figure 7A) and did not induce significant changes in cell death when SNX631 was added at the first exposure to suspension (P1 DMSO vs SNX631, Figure 7B), we tested if pretreatment of attached cells with SNX631 followed by suspension culture for 24 hours, with or without SNX631, would affect cell death. We find that pretreatment of SNX631 in 2D reduced the survival of adapted OV90 AnR cells in suspension to the level of their AnS parent cells (Figure 7D, 2D SNX-suspension DMSO condition). Pretreatment with SNX631 (2D) followed by maintaining the treatment in suspension culture further increased the cell death of adapted OV90 AnR cells in suspension by as much as 40% (Figure 7D, 2D SNX-suspension SNX condition). In contrast, pretreatment (2D) of parental OV90 AnS cells with SNX631 did not significantly increase cell death in suspension (Figure 7D, 2D SNX-suspension DMSO condition). However, when SNX631 was also present during suspension culture, AnS cells exhibited 25% more cell death (Figure 7D, 2D SNX-suspension SNX condition). Strikingly, vehicle pretreated AnR cell (2D) with SNX631 added only in suspension exhibited significantly higher cell death (Figure 7D 2D DMSO, suspension SNX condition) as well, as compared to parental AnS cells under the same conditions which exhibited no significant change in cell death (Figure 7D, 2D DMSO-suspension SNX condition and seen in Figure 7B, P1). These data together suggest that SNX631 promotes anoikis sensitivity with a heightened vulnerability of AnR cells to such CDK8/19 inhibition, and demonstrates that CDK8/19 inhibition can both prevent and reverse AnR.

### CDK8/19 inhibition disrupts the balanced transcriptional reprogramming underlying both the development of anoikis resistance and the acute response to anoikis stress

To investigate the transcriptional basis for CDK8/19 inhibition’s effects on restoring anoikis sensitivity in anoikis-resistant cells (AnR) we performed RNA-Seq analyses on both anoikis-sensitive (AnS) and adapted anoikis-resistant (AnR) OV90 cells following 4 days of SNX631 treatment under standard 2D culture conditions. Since AnR cells showed enhanced anoikis susceptibility to SNX631 (Figure 7D) we also profiled AnR cells under suspension culture conditions, with or without SNX631 treatment, to capture the transcriptional response of AnR cells to anoikis stress (schematic, Figure. 8A).

**Figure 8.**
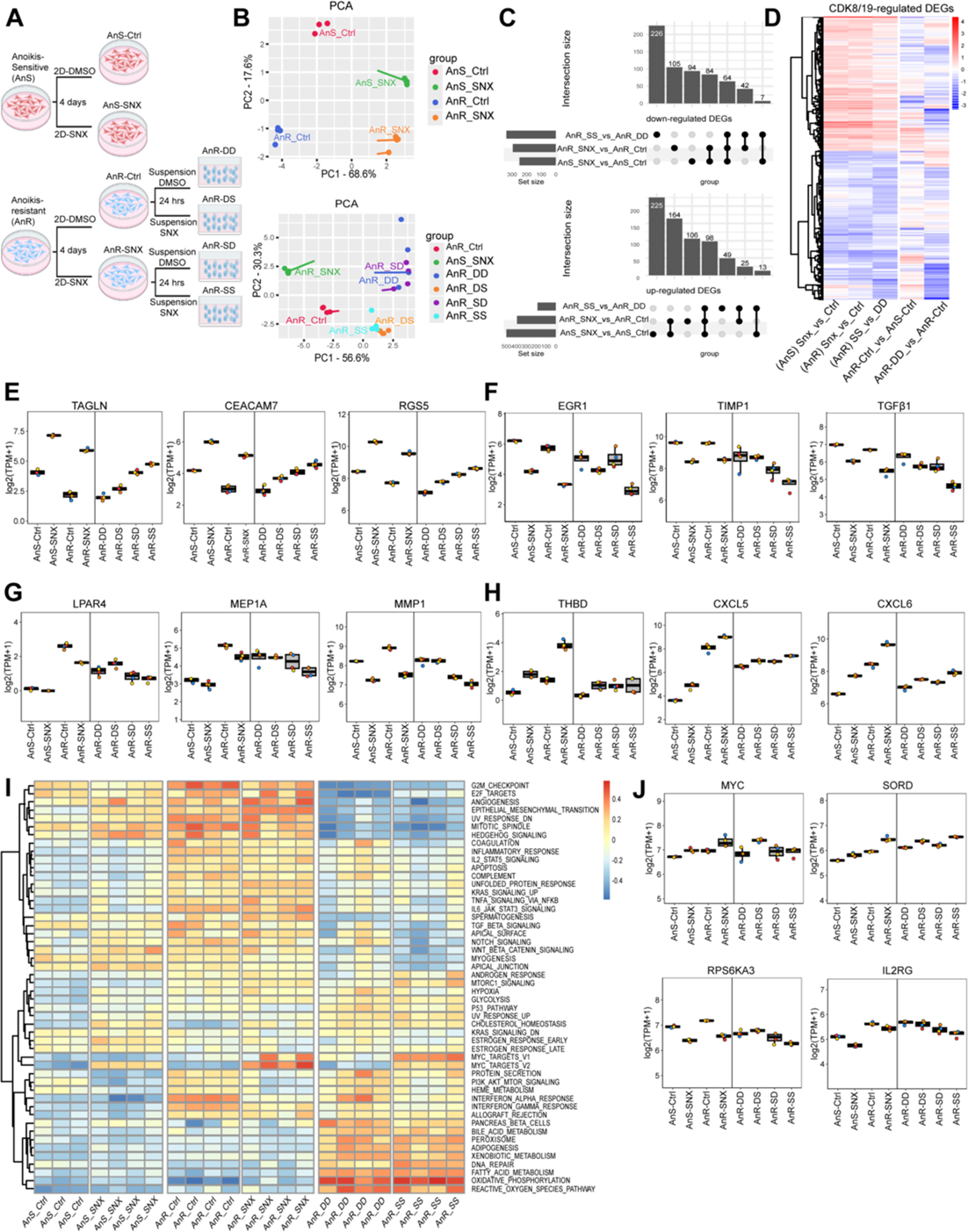
CDK8/19 inhibition induces bidirectional transcriptional reprogramming in anoikis-sensitive and -resistant cells. **(A)** Schematic of the RNA-Seq experimental design. OV90 anoikis-sensitive (AnS) and anoikis-resistant (AnR) cells were treated with either DMSO or SNX631 for 4 days under standard 2D culture conditions. AnR cells pre-treated in 2D conditions were then further cultured in suspension, with DMSO or SNX631. **(B)** Principal component analysis (PCA) of transcriptomic profiles: AnS vs. AnR cells under 2D culture (Top), and AnR cells under 2D vs. suspension conditions (Bottom). **(C)** UpSet plot showing intersections of differentially expressed genes (DEGs) regulated by SNX631 in three comparisons: AnS cells in 2D (AnS-SNX vs. AnS-Ctrl), AnR cells in 2D (AnR-SNX vs. AnR-Ctrl), and AnR cells in suspension (AnR-SS vs. AnR-DD). DEGs were defined by FDR < 0.01 and fold change > 1.5. **(D)** Heatmap displaying the union of all SNX631-regulated DEGs across the comparisons in (C). Columns 1–3: transcriptional response to SNX631 under different conditions; Column 4: DEGs associated with development of anoikis resistance (AnR-Ctrl vs. AnS-Ctrl); Column 5: DEGs associated with acute anoikis response in AnR cells (AnR-DD vs. AnR-Ctrl). **(E, F)** Expression levels (TPM) of representative DEGs that are upregulated **(E)** or downregulated **(F)** by CDK8/19 inhibition in AnR cells under suspension conditions (AnR-SS vs. AnR-DD). **(G, H)** Expression levels of representative genes showing bidirectional regulation by CDK8/19 inhibition in AnR cells: reversed **(G)** or enhanced **(H)** expression compared to untreated conditions. **(I)** GSVA heatmap of hallmark pathway enrichment across individual samples. Pathways are clustered based on Euclidean distance using normalized enrichment scores (NES). **(J)** Expression levels of representative genes from the MYC Targets V1 and PI3K/AKT/mTOR signaling hallmark pathways.

Principal component analysis (PCA) of the RNA-Seq data revealed a clear separation between AnS and AnR cells along both PC1 (68.6% variance) and PC2 (17.6% variance), with SNX631 treatment shifting the transcriptomes of both cell types in a similar direction. Notably, the transcriptional changes induced by CDK8/19 inhibition on PC1 were opposite to those associated with the development of anoikis resistance (Fig. 8B, top). In AnR cells, transitioning from 2D to suspension culture (AnR-DD vs. AnR-Ctrl) also led to a pronounced shift along PC1 (56.6% variance), consistent with transcriptional changes in response to anoikis stress. This shift was partially reversed by SNX631 treatment in 3D culture, in the same direction as the effect of CDK8/19 inhibition under 2D conditions (Fig. 8B, bottom).

Differential expression analysis (cutoffs: fold change >1.5, FDR <0.01) identified 749 DEGs (500 upregulated, 249 downregulated, Supplemental Table 1) in AnS cells treated in 2D, 688 DEGs (393 up, 295 down, Supplemental Table 2) in AnR cells treated in 2D, and 524 DEGs (185 up, 339 down, Supplemental Table 3) in AnR cells treated in 3D. Upset analysis (Figure. 8C) showed that 497 genes were shared between at least two of these conditions, suggesting that CDK8/19 regulates a core set of genes regardless of culture conditions or anoikis resistance status. Heatmap analysis of the union of all SNX631-regulated DEGs (from all three comparisons) further highlighted the consistent transcriptional response induced by CDK8/19 inhibition across the different OV90 cell states (Fig. 8D). Remarkably, many of these SNX631-responsive genes were also altered during the acquisition/development of anoikis resistance (AnR-Ctrl vs. AnS-Ctrl) or upon exposure to anoikis stress (AnR-DD vs. AnR-Ctrl), suggesting that CDK8/19 activity is essential for maintaining the balanced transcriptional programs that enable aggressive ovarian cancer cells to survive under detachment stress.

Examination of a subset of the top DEGs upregulated or downregulated by CDK8/19 inhibition in AnR cells under suspension conditions (AnR-SS vs. AnR-DD, Figure 8E,F) reveal upregulated genes including *TAGLN, CEACAM7*, and *RGS5*, while downregulated genes include *EGR1,* previously identified as a SNX631-responsive gene in other cellular models [33, 49, 55, 56], as well as *TIMP1* and *TGFβ*1, also known for its role in anoikis resistance and ovarian cancer metastasis. [23, 57–59]. Notably, most of these genes are downregulated in AnR cells, with some further decreased upon suspension culture. We also examined DEGs upregulated in AnR cells and found two distinct responses to CDK8/19 inhibition: induction of some genes was reversed (e.g., *LPAR4, MEP1A, MMP1*; Figure. 8G), whereas others were further enhanced (e.g., *THBD, CXCL5, CXCL6*; Figure. 8H). Additionally, we observed that omitting SNX631 treatment from either 2D (AnR-DS) or suspension conditions (AnR-SD) diminished the transcriptional regulatory effects of CDK8/19 inhibition on most of these genes, consistent with the most significant effects on anoikis observed upon continuous treatment (Figure 7C) and suggests that sustained CDK8/19 activity is required for maintaining the dynamic transcriptional programs underlying the transcriptional plasticity. GSVA analysis using Hallmark gene sets (Figure. 8I) revealed widespread pathway alterations across different cell states. Among these, several pathways upregulated in AnR cells were suppressed by CDK8/19 inhibition, including PI3K/AKT/mTOR signaling, heme metabolism, and interferon responses. Conversely, the MYC targets gene set, upregulated in AnR cells, were further increased by SNX631, indicating that CDK8/19 inhibition induces bidirectional disruption of transcriptional pathways. Representative genes illustrating these effects include *RPS6KA3* and *IL2RG* (PI3K/AKT/mTOR pathway) and

*MYC* and *SORD* (MYC pathway) respectively (Figure. 8J). Together, these analyses demonstrate that CDK8/19 inhibition disrupts balanced, pathway-specific transcriptional reprogramming that underlies key mechanisms of anoikis resistance.

## DISCUSSION

We report here that OC cells exhibit a range of sensitivity to cell death upon matrix detachment in suspension (anoikis). Cell line models that exhibit AnS, when subjected to cycles of detachment and reattachment in vitro, acquire AnR. Such acquired AnR mimics the AnR that develops after intraperitoneal ovarian cancer growth and tumor cell survival in ascites, but is not permanent, as cells can revert to their baseline state in the absence of detachment stress. Importantly, the acquired adaptive anoikis resistance is associated with transcriptional reprogramming and increased migratory capabilities, changes in mitochondrial capacity, resistance to chemotherapy, ability of cells to metastasize to the lungs and evasion of immune cell killing. Critically, selective inhibition of CDK8/19 Mediator kinase, a regulator of transcriptional reprogramming, prevents and suppress anoikis adaptation and metastasis in OC models while rebalancing the transcriptional program.

Ovarian cancer metastasis requires cell detachment from primary sites and survival in peritoneal cavity or circulation upon loss of attachment. Anoikis-resistant cells in ascites and circulation correlate with poor outcomes, relapse, and chemoresistance.[2, 4, 8, 12, 15, 60–66] Notably, whole exome sequencing has suggested that recurrent ascites tumors contain mutations already present initially [67] indicating epigenetic/transcriptional adaptation rather than genetic evolution drives recurrence. To model, this we exposed AnS OC cell lines to intermittent (24 hours) detachment induced stress followed by re-attachment intervals. These attachment-detachment cycles gradually improved survival under suspension, through reduced apoptosis rather than enhanced proliferation or clonal selection. The survival capacity of individual clones fluctuated over time, indicating ability to interchange between sensitive and resistant states, suggesting non-mutational adaptation, which was confirmed by whole exome sequencing. Our reversion experiments without detachment stress revealed that the acquired resistance was not permanent and non-genetic in nature. The findings herein were not concordant with two states and suggest likely additional states in the population. The Coefficient of Variation (CV) analysis implies the existence of either pre-established cellular memory or multiple cellular states within the population. This diversity results in varied responses to detachment stress, with individual clones showing greater behavioral differences than control populations. Such cellular heterogeneity warrants future investigation for understanding stress adaptation mechanisms.

A central question that we pursued was whether a non-permanent anoikis resistant phenotype was sufficient for metastatic capabilities. Indeed, adapted anoikis resistant cells were more motile, consistent with prior studies showing concomitant increases in migration and anoikis resistance [68], phenotypes that are reciprocally related as forced confined migration can also induce anoikis resistance. [44] Chemosensitivity changes while consistent across several cell lines appeared specific to paclitaxel. Although cisplatin and doxorubicin sensitivity appeared unchanged, testing under conditions that mimic anoikis selection (brief drug exposure followed by regrowth assessment) may reveal additional resistance mechanisms not captured by the standard viability assays. The adapted anoikis resistant cells also showed increased oxygen consumption, ATP production, and spare respiratory capacity, likely enabling functional maintenance under detachment stress. AnR cells also exhibited elevated extracellular acidification rates (ECAR), indicating simultaneous upregulation of glycolysis alongside OXPHOS. This metabolic flexibility of high OCR and ECAR could create therapeutic opportunities through dual metabolic targeting that combines OXPHOS inhibitors (which AnR cells showed heightened sensitivity to) with glycolytic inhibitors (targeting LDH or MCT) or pH regulation disruptors (CAIX inhibitors), which could eliminate compensatory metabolic escape and selectively stress these adaptable cells. Beyond metabolic changes, AnR cells showed reduced susceptibility to T cell-mediated killing, demonstrating functional immune evasion, and reversing these phenotypes could also contribute to improvements in immune checkpoint therapies, that currently have limited use in the management of ovarian cancer. These multiple adaptations, metabolic flexibility, immune escape, and chemoresistance, highlight why preventing AnR development and reversing specific genes and pathways, may be more effective than targeting a specific resistance mechanism. Our finding that CDK8/19 inhibition could accomplish both, suggests this as a more tractable therapeutic strategy than targeting a single pathway of established resistance.

Moreover, while the acquisition of metastatic phenotypes in the anoikis adapted cells and changes in hallmark pathways during anoikis adaptation were seen across ovarian cancer models, the timing of distinct pathways was somewhat altered in each model. CAOV3 cells are significantly more sensitive to matrix detachment stress compared to OV90 cells, carry different mutations (depmap.org) and most notably revert/lose their acquired resistance sooner than OV90. Thus it remains to be determined whether the inherent differences in baseline sensitivity to matrix detachment stress and/or the mutational backgrounds influenced the timing differences observed. This variability could present a therapeutic challenge, as manifested in the scarcity of targeted treatments universally applicable to the clinical management of ovarian cancer. Regardless of the timing and some pathway differences seen between models, the dependency of anoikis adaptation on transcriptional reprogramming was evident in marked sensitivity to CDK8/19 Mediator kinase inhibition across cell lines. CDK8 and CDK19, are alternative enzymatic components of the CDK module that regulates the transcriptional Mediator complex and are not essential to the overall transcription machinery. [69] Consistent with this selective role, the CDK8/19 inhibitor SNX631 had minimal effects on steady-state growth of ovarian cancer cells at 500 nM, a concentration sufficient for complete kinase inhibition[70], yet potently blocked the development of anoikis resistance across multiple ovarian cancer models. This selective vulnerability aligns with prior studies showing CDK8/19 inhibition prevents adaptive drug resistance. [49, 71] Strikingly, SNX631 not only prevented anoikis resistance development but also resensitized anoikis-resistant cells, with maximal effects achieved through combined pretreatment and acute exposure during suspension stress. Our transcriptomic analysis revealed that this resensitization involves complex transcriptional reprogramming rather than simple phenotypic reversal, as the DEGs between SNX631-treated AnR cells and untreated AnS cells exceeded the sum of individual effects, indicating that CDK8/19 inhibition creates distinct transcriptional states while disrupting critical resistance nodes. CDK8/19 inhibition induced bidirectional transcriptional effects, with approximately 50% of changes shared between AnS and AnR cells representing a core CDK8/19-dependent program. Some resistance pathways were reversed while others were paradoxically enhanced. This dual pattern reflects CDK8/19’s established role as both a positive and negative transcriptional regulator, [33] functioning as a molecular rheostat that maintains the transcriptional balance required for anoikis resistance. Of note, RNA levels of CDK8/19 were not changed during adaptation, hence the heightened vulnerability of AnR cells could also reflect transcriptional addiction to a CDK8/19 mediated stress response programs with the selective disruption of this balance, rather than global suppression, leading to increased sensitivity. The persistent dependency on CDK8/19 activity even under active suspension conditions where anoikis resistance is functionally critical, suggests that repeated detachment stress creates lasting reliance on these transcriptional mechanisms.

The selective targeting of stress-induced transcriptional plasticity while sparing basal transcription makes CDK8/19 an attractive therapeutic target for preventing metastatic adaptation. Although CDK8/19 expression is elevated in several ovarian cancer subtypes and correlates with patient survival, [36, 56, 72] our findings suggest that the therapeutic efficacy of CDK8/19 inhibition depends on targeting transcriptional dependencies rather than expression levels alone. Furthermore, genes upregulated in anoikis resistance (Figure 8) that are reversed by SNX631 could serve as biomarkers to identify patients most likely to benefit from CDK8/19 inhibition. Given ovarian cancer’s extraordinary adaptability during intraperitoneal dissemination and limited targeted therapy options, CDK8/19 represents a novel vulnerability where transcriptional dependencies could guide both patient selection and therapeutic intervention.

## MATERIALS AND METHODS

### Cell lines and culture conditions

Cell lines were cultured in their basal medium, 10% FBS, and 100 IU/ml penicillin 100µg/ml streptomycin unless indicated in Table 1 and were maintained in a 5% CO2 incubator at 37^°^C. Cell line authentication was done using STR profiling at the UAB Heflin Center for Genomic Sciences. Cells were tested frequently for mycoplasma contamination using LookOut® Mycoplasma PCR Detection Kit (cat no: MP0035-1KT, Millipore sigma). CAOV3, P76, P151, P201, P211, HEK293 were cultured in DMEM (contains high glucose, L-glutamine and sodium pyruvate) supplemented with 10% fetal bovine serum (FBS). TYK-nu was cultured in DMEM+1% MEM non-essential amino acids+1% MEM vitamins. ID8-EMD, ID8-BRCA2, ID8-TP53 -/- were cultured in DMEM 4% FBS, 5 µg/mL insulin, 5 µg/mL transferrin and 5 ng/mL sodium selenite. FT282 was cultured in 1:1 mixture of DMEM and F12 medium supplemented with 10% FBS and 2mM L-glutamine. OV90, EOC15 and IOSE141 were cultured in 1:1 mixture of MCDB 105 and Medium 199 supplemented with 15% and 10% FBS respectively. SK-OV3, OVCA433, OVCA420, HEY, HEYA8 and OVCAR10 were cultured in RPMI (contains l-glutamine) supplemented with 10% FBS and OVCAR3 with 20% FBS. OVCAR4 and OVCAR5 were cultured in RPMI supplemented with 10%FBS, 2mM glutamine, 0.25 U/mL insulin. All additional resources are listed in Tables (3-8).

**Table 1:** Cell density for IC50 experiments.

| Cell line | Initial cell density |
| --- | --- |
| OV90 | 5000 |
| OVCA420 | 2000 |
| CAOV3 | 5000 |
| EOC15 | 5000 |

**Table 2:** Cell density for transwell fibronectin migration assay.

| Cell line | Initial cell density | Incubation time (hours) |
| --- | --- | --- |
| OV90 | 100000 | 24 |
| CAOV3 | 100000 | 24 |
| OVCA420 | 75000 | 24 |
| EOC15 | 100000 | 24 |
| P151 | 75000 | 6 |

**Table 3.**
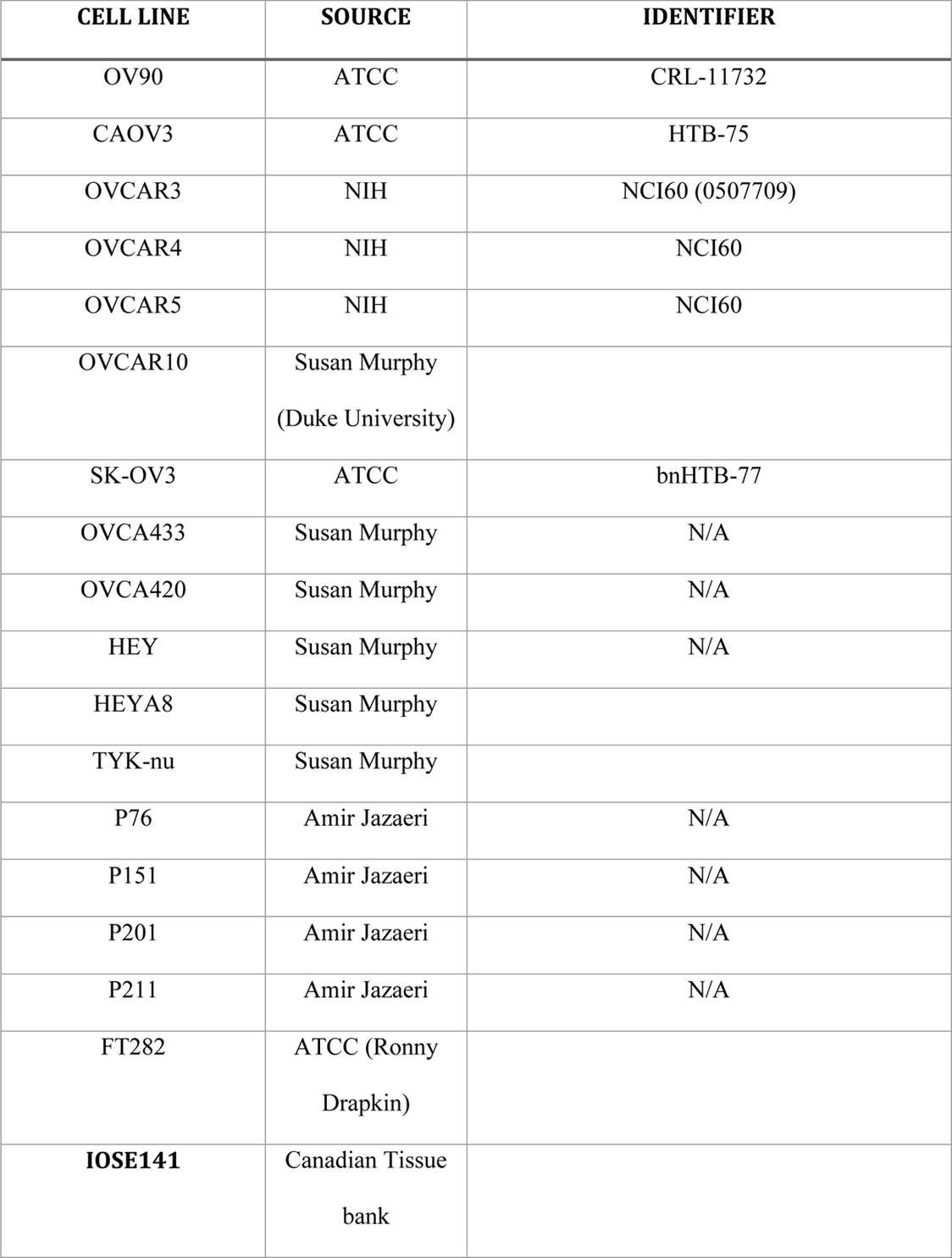
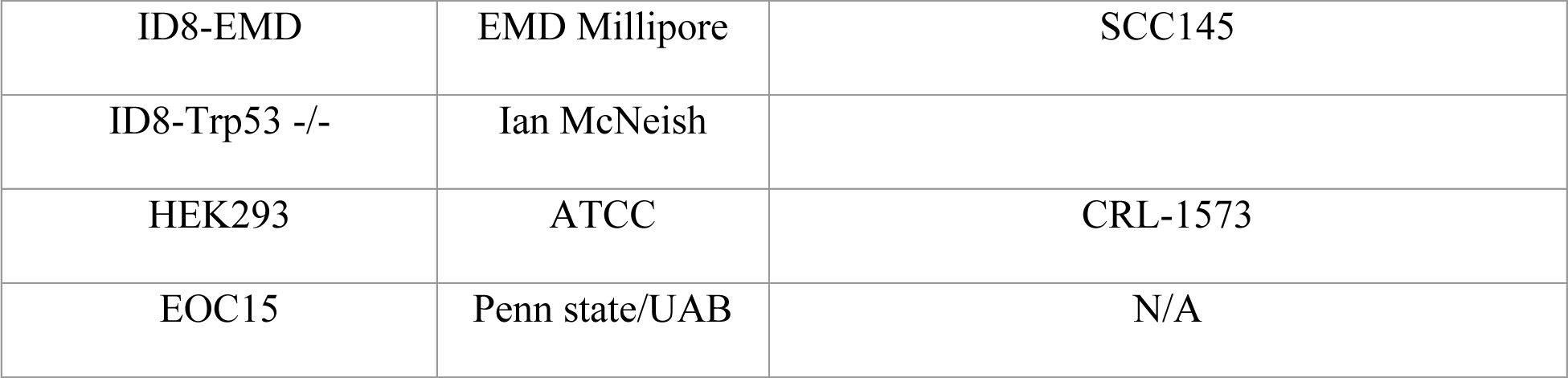
Cell lines.

**Table 4:** Commercial Kits.

| Reagents | Source | Catalog number |
| --- | --- | --- |
| LIVE/DEAD™<br>Viability/Cytotoxic kit | Fisher Scientific | L3224 |
| LookOut® Mycoplasma<br>PCR Detection Kit | Millipore sigma | MP0035-1KT |
| Annexin V Apoptosis<br>Detection Kits | eBioscience™ | 88-8005-74 |
| CD8+ T cells isolation<br>kit | Miltenyi Biotec | 130-096-495 |
| DNeasy Blood and<br>Tissue Kit | Qiagen | 69504 |

**Table 5:** Antibodies.

| **Antibodies** | **Source** | **Catalog number** |
| --- | --- | --- |
| Ki-67 (8D5) Mouse<br>mAb | Cell signaling | 9449 |
| Goat anti-Mouse IgG1<br>Cross-Adsorbed<br>Secondary antibody<br>Alexa Fluor™ | Invitrogen | A-21125 |
| Cleaved caspase 3 | Cell signaling<br>technology | 9661T |
| YH2AX | Cell signaling<br>technology | 9718T |
| annexin V | Bio Legend | 640906 |
| PI | Invitrogen | 00-6990-42 |
| pSTAT1 | Cell signaling<br>technology | 9177S |
| MHC1 | R&D | NBP3-16696 |
| TAP1 | Cell signaling<br>technology | 49671S |
| MYC | Cell signaling<br>technology | 13987 |

**Table 6:**
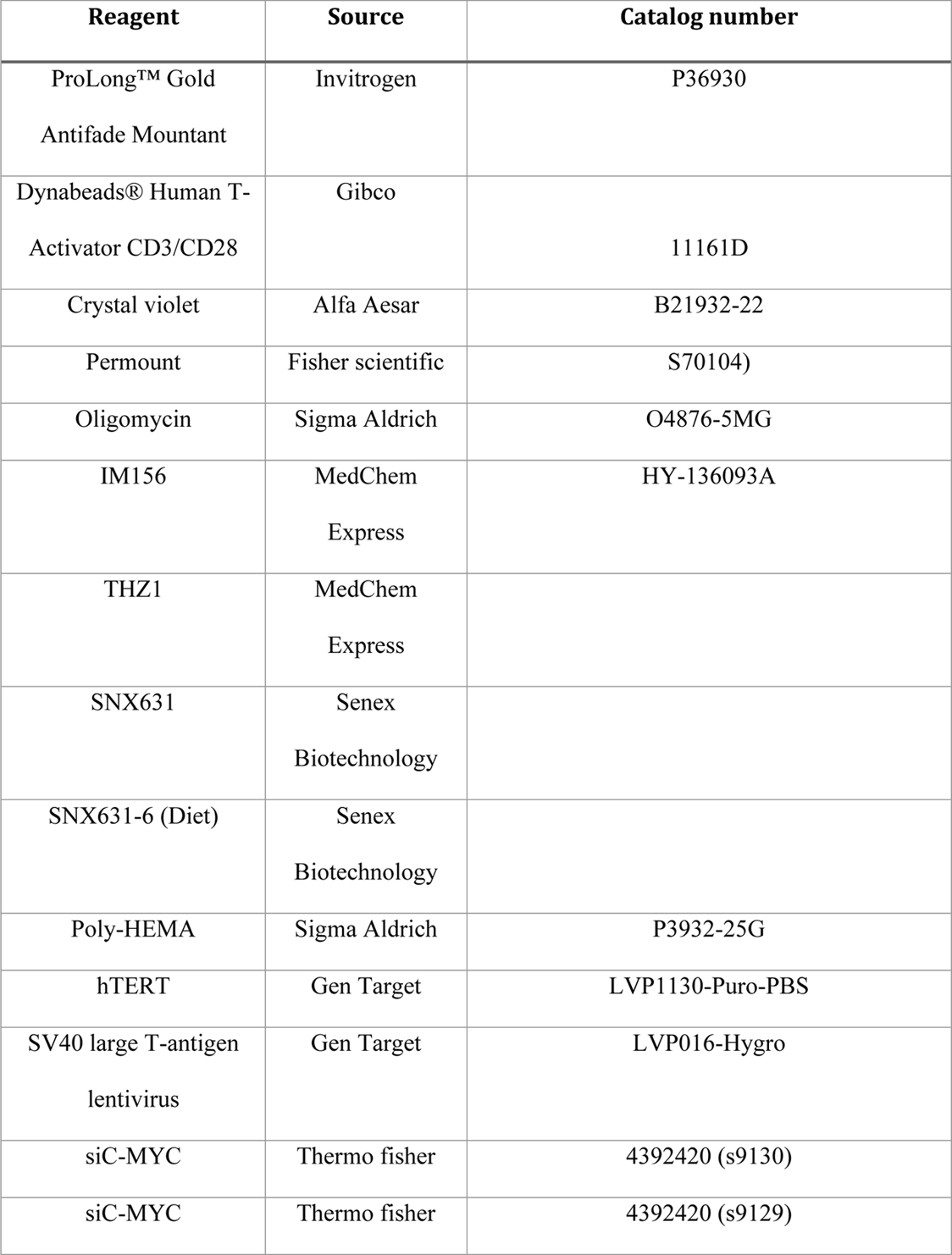
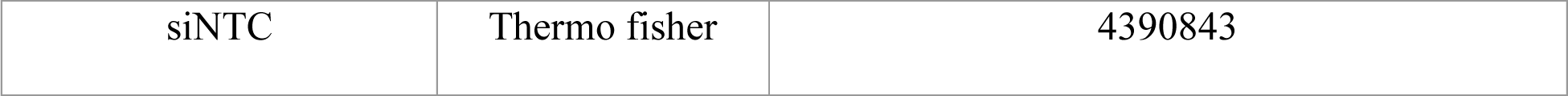
Other reagents and recombinant constructs.

**Table 7:** Software used.

| Software | Source | Identifier |
| --- | --- | --- |
| ImageJ | NIH | ImageJ,<br>RRID:SCR_003070 |
| Prism9 | GraphPad | GraphPad Prism<br>(RRID:SCR_002798)( |
| Adobe Illustrator | Adobe | Adobe Illustrator,<br>RRID:SCR_010279 |
| **FlowJo** |  | FlowJo (RRID:SCR_008520) |

**Table 8.**
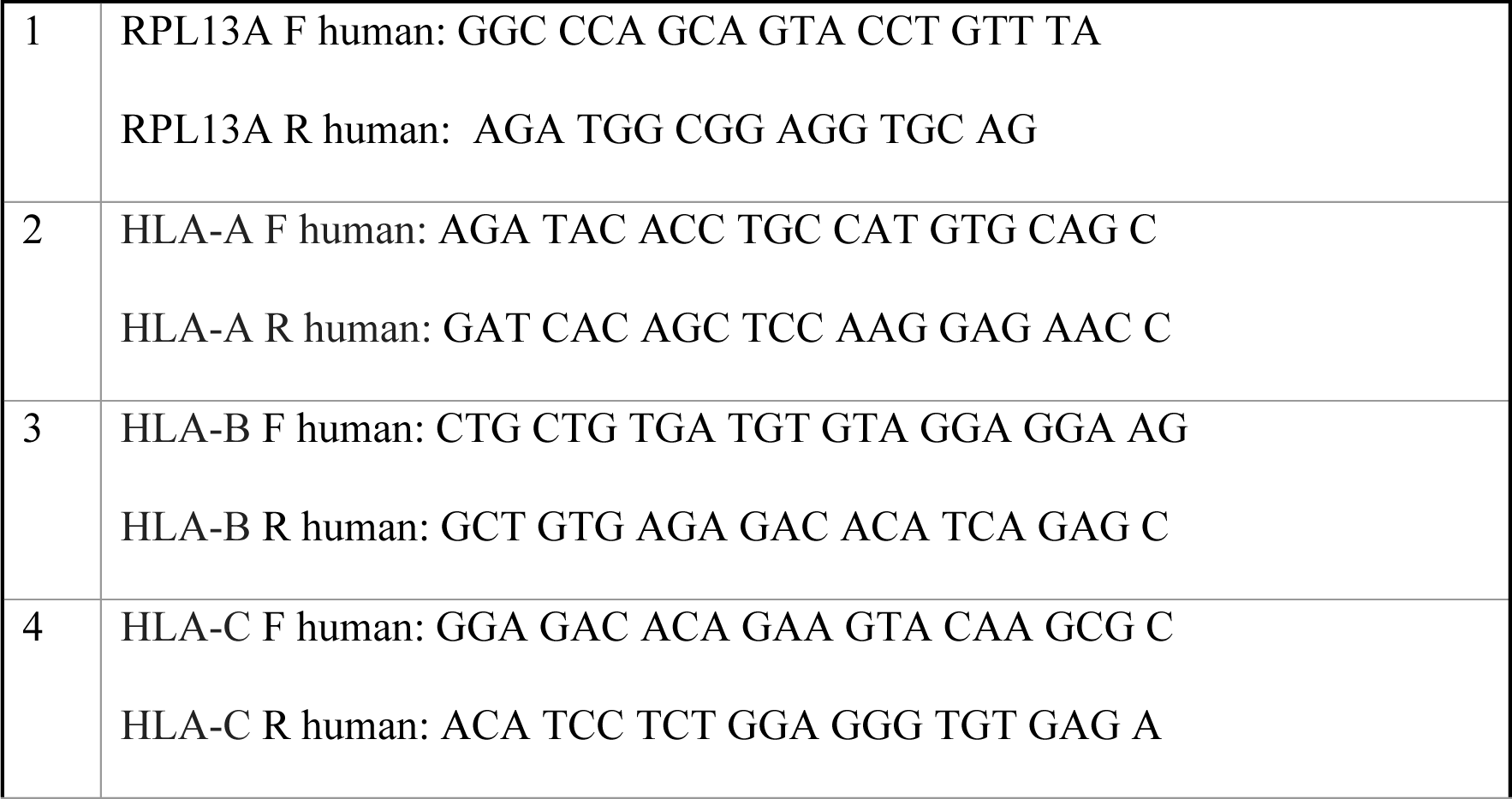
qRT-PCR Primers (listed 5’ to 3’)

### Live cell/viability assessment of cell aggregates in suspension (multiple spheroids or cell aggregates)

6 well plates were coated with poly-HEMA (2% in 95% ethanol), then 1 ml poly-HEMA was added to each well and allowed to dry at 50° C overnight. The plates were UV-radiated for 1 hour. At this point, the plates were used for experiments or wrapped with Parafilm and kept at 4°C. Cells were cultured in 100mm plates until they reached ∼90% confluency then trypsinized and counted with Trypan blue using Countess II FL Automated Cell Counter. 250,000 cells were cultured in poly-HEMA coated 6 well plates in full serum under regular growth medium (3 ml per well) for 24 hours. The cells from each well were collected in a 15 ml tube and centrifuged at 1200 rpm for 5 minutes. Next, the medium was removed and 100 µl of 10x Trypsin-EDTA (T4174-100ML Millipore sigma) was added to the cells and incubated for 5-20 minutes (depending on cell line and spheroid compactness) in the incubator. 400 µl medium was added to the cell suspension and the number of live cells was counted with Trypan blue exclusion using Countess II FL Automated Cell Counter.

### Live cell/viability assessment of single spheroids in suspension (single spheroid)

To obtain single spheroids, 1,000 OV90 and 2,500 CAOV3 cells were transferred to Ultra low attachment U bottom black wall 96 well plates (Corning: 4515) in 200 ul medium. The plate was centrifuged at 1200 rpm for 2 minutes and then incubated for 24 hours. For live dead assays, the LIVE/DEAD™ Viability/Cytotoxicity Kit, for mammalian cell: L3224 was used. Briefly, green-fluorescent Calcein-A and red-fluorescent ethidium homodimer-1 were added to HBSS buffer (Hanks’ Balanced Salt Solution) or PBS at 4 and 8 µM respectively. Then 100 µl of the medium was removed from each well and 100 µl HBSS containing dyes were added slowly to each well to have a final concentration of 2 and 4 µM of Calcein and Ethidium. After incubating the cells for 90 minutes, washing was done 3 times very carefully to avoid disrupting the spheroids (100µl of the dye solution was removed, and 100 µl of HBSS was added). Imaging was done using a confocal microscope (Nikon TE2000 inverted) at UAB’s High Resolution Imaging Facility and images were analyzed in Image J.

### Cyclic cell culture

250,000 cells were cultured in poly-HEMA coated 6 well plates for 24 hours. Cell number and dish size were kept constant to obtain reproducible live cell numbers. After 24 hrs., the cells were centrifuged at 1200 rpm for 5 minutes and viability was measured as described under ‘ *Viability assessment of cell aggregates in suspension’* prior to proceeding to plating the cells into standard tissue culture 2D conditions in a 60mm TC dish until they reached 90% confluency. The media was changed every 3-4 days. The first three steps were repeated for another 6-8 cycles depending on the cell line (total 7-9 passages).

### Memory/reversion studies

Adapted anoikis resistant (AnR) OV90 and CAOV3 cells from suspension culture were seeded into 60mm dishes and sub cultured in 2D for a total of 9-11 passages in 60mm dishes. At each passage, when the cells became 80-90% confluent, a fraction of cells were cultured in suspension in polyhema-coated plates for 24 hours and live cell/viability was measured as described above under ‘*Viability assessment of cell aggregates in suspension’*. The remaining cells were maintained in 2D. The number of generations for these experiments was calculated as the number of times the cell population doubled over time. The number of generations in each passage was calculated using: log2 (𝑥) where x is equal to the (final cell number/initial cell number.)

### Doubling time (DT) measurement in 2D

5000 cells were cultured in a 96 well plate in 100 µl medium and SRB assay as described previously [73] and was done every 24-48 hours for 7 days. Absorbance was measured at 570 nm using a Synergy H1 microplate reader (Biotek). DT was calculated using absorbance values obtained from the growth phase of the cells using the equation: 𝐷𝑇 = 𝑙𝑛2/((𝑙𝑛 [𝑂𝐷2/𝑂𝐷1 ]) / (𝑇2 − 𝑇1)). OD1 and OD2 are the absorbance values measured in Time 1 and 2 (T1, T2) respectively.

### Doubling time measurement in suspension

250,000 cells were cultured in suspension in a 6 well poly-HEMA coated plate. Cell viability was measured using a Trypan blue exclusion assay every 24-48 hours for 10 days. Doubling time (DT) was calculated using the equation: DT = ln2/((ln [CN1/CN2 ]) / (T2-T1)). CN1 and CN2 are the cell number values measured in Time 1 and 2 (T1, T2) respectively.

### Flow cytometry-based apoptosis assay

Parental and AnR OV90 and CAOV3 cells were cultured in suspension for 24 hours as described above. The cells were treated with Trypsin –EDTA for 5-10 minutes to produce a single-cell suspension. Media was added to the cell suspension to neutralize trypsin. Then, 500,000-100,0000 cells were transferred to a new 1.5 microcentrifuge tube and washed with Cold PBS twice. The cells were stained with Annexin V and PI using according to the manufacturer’s protocol. Flow cytometry was done using BD LSRFortessa (Europa) at the UAB flow cytometry core. Analysis was carried out in FlowJo 10.8.1.

### Ki67 Immunofluorescence staining in suspension

Suspension cells (1×10^6^) were collected after 24h and centrifuged (1200 rpm, 5min). Cells were washed in cold PBS, fixed (4% PFA, pH 7.4, 15min), quenched (10mM NH4Cl, 5min), permeabilized (0.3% Triton X-100, 10min), and blocked (5% BSA, 1h). Cells were incubated with Ki-67 antibody (1:450 in 3% BSA, overnight, 4°C), followed by Alexa Fluor conjugated secondary antibody incubation (1:500, 1h) and DAPI (1:2000, 10min). After cytospinning (800 rpm, 5min), slides were mounted with ProLong Gold. Images were captured using Nikon TE2000 confocal microscope and analyzed in ImageJ. Ki-67 CTCF (integrated density minus area × background) was normalized to DAPI CTCF

### 3D Cell viability assessment in single clones

Single Cell cloning was carried out by serial dilution of OV90 cells in 96 well plates (Corning protocol) (https://www.corning.com/catalog/cls/documents/protocols/Single_cell_cloning_protocol.pdf). After 2 weeks, single clones were trypsinized and seeded in 6 well plates and passaged in attached conditions (2D) for 6 passages. The viability of each clone in suspension from passages 1,3, and 6 was analyzed using Trypan blue exclusion.

### Patient and mouse ascites derived cells

Patient ascites cells were established as previously described [19, 23] with prior approval from the Penn State College of Medicine. Patient-derived (50 ml) or mouse ascites fluid (post-OV90/ID8 i.p. injection) was centrifuged (4000 rpm, 20 min). RBC lysis buffer was added to the pellet (10:1 v/v, 10 min RT), followed by PBS addition and centrifugation (300g, 10 min). Cells were resuspended in media and counted using Countess II FL and cultured in poly-HEMA plates (250k cells/well, 2 weeks), disaggregated after one week, and maintained in 2D. For immortalization, cells (50,000/well) were transduced with hTERT and SV40 large T-antigen lentivirus (MOI 15) with polybrene (10 µg/ml) for 24h, followed by 48h recovery and puromycin selection (5 µg/ml, 48h)

### Immune mediated tumor killing assay

CD8+ T cells were isolated from healthy PBMCs (Precision for Medicine) using Miltenyi CD8+ isolation kit (#130-096-495). T cells were cultured (500,000 cells/well) in RPMI+10% FBS+30U/ml IL2. For activation, T cells (1×10^6) were incubated with CD3/CD28 Dynabeads (25µL) for 48h. Tumor cells (10,000 OV90 or CAOV3; parental, P1, or P7) were seeded 24h before adding activated T cells at 10:1 ratio. Surviving tumor cells were counted at 24h and 48h using trypan blue exclusion after washing. For apoptosis assessment, tumor cells were stained with cleaved caspase-3 antibody (1:100) after T cell co-culture (12h for CAOV3, 48h for OV90), and CC3-positive cells were quantified using ImageJ

### SNX631 and THZ1 treatment in OV90 cells, OVCA420

The cells were cultured in TC-treated plates until 80% confluent. Following trypsinization, 250,000 cells were cultured in poly-HEMA coated 6 well plates in medium containing either DMSO, or indicated concentrations of SNX631 or THZ1. After 24 hours of incubation, live cell count/viability was measured using Trypan blue, and 500,000 cells were seeded in 60mm dishes in the presence of either DMSO, SNX631, or THZ1 until 80% confluent. This cycle of attachment-detachment (in the presence of drugs) continued as described in the figures.

### SNX631 Pre-treatment

800,000 OV90 parental or AnR cells were seeded in 60mm dishes in the presence of either DMSO or 500nM SNX631 for 96 hrs. Cells were trypsinized and 250,000 cells were cultured in poly-HEMA coated 6 well plates in a medium containing either DMSO or SNX631 for 24 hrs. Live count/ viability was measured using trypan blue.

### Oligomycin and IM156 treatment in OV90 and CAOV3 in suspension culture

250,000 OV90 or CAOV3 from either P0 (Parental) or P7 (undergone 7 cycles of attachment-detachment) were cultured in poly-HEMA coated 6 well plates in their respective growth media containing DMSO as the control and either 1.5 µM Oligomycin or IM156 at 20 µM for CAOV3, 15 and 25 µM respectively for OV90 cells and incubated for 72 hours. Next, the cells were collected and trypsinized in 10x Trypsin for 10 minutes to make single-cell suspensions, and cell death and apoptosis were measured by flow using annexin/PI staining.

### Seahorse Mitochondrial stress test

Seahorse Mito stress test was done following the UAB Bioanalytical Redox Biology (BARB) core XF96 assay protocol. Specifically, OV90 (20,000) and CAOV3 (25,000) cells (P0, P1, and P7) were cultured in XF96 plates for 36h. The XF96 cartridge was hydrated overnight, then loaded with calibrant (1h, 37°C, non-CO2). Cells were washed and incubated in XF media (DMEM with 10mM glucose, 2mM glutamine, 1mM pyruvate) for 1h. Ports were loaded with: A) oligomycin (1.5µM final), B) FCCP (0.675 or 0.9µM), C) FCCP (1.65 or 1.8µM), and D) antimycin A/rotenone (0.5µM each). The assay cycle (3min mix, 3min measure) was performed after each port injection. Analysis used Seahorse Wave 2.6 and GraphPad Prism 10.0.2.

### IC50 determination

Ovarian cancer cells were seeded in 96 well plates at indicated cell density/well (Table 1) and incubated until 50-60 % confluent. Cell growth medium containing drugs (Cisplatin, Doxorubicin, Paclitaxel, IM156, SNX631, or THZ1) was added in 8 dilutions to the cells and incubated for 48 hours to 7 days followed by SRB assay. Absorbance was measured at 570 nm using a Synergy H1 microplate reader (Biotek). IC50 was calculated using nonlinear regression (four parameters) in GraphPad Prism software 10.0.2.

### Transwell fibronectin migration

24 well transwell with 8 μm pore membrane (Greiner bio-one, catalog: 662638) was coated with 10 μg/ml fibronectin in sterile DI water solution and incubated at 37° C for 2 hours. Then cells were suspended in 100 µl serum-free medium and added to the upper chamber of each transwell (refer to Table 2 for cell number and incubation time). The lower chamber was filled with 600 µl Full serum medium and it was incubated in a CO2 incubator at 37° C for indicated hours. Then the medium was removed and transwells were washed with PBS twice. Non-invaded cells were scraped by a cotton swab. Then cells were fixed with 4% PFA (paraformaldehyde, Avantor, catalog: S898-07) at RT for 5 minutes. Cells were permeabilized with 100 % ethanol for 20 minutes. Next, the cells were stained with 0.5% Crystal violet (Alfa Aesar catalog: B21932-22) for 15 minutes and washed with DI water. Transwells were air dried at RT overnight and membranes were cut and mounted on glass slides with Permount (Fisher scientific catalog: S70104). Imaging was done using EVOS M7000 inverted microscope (Thermo fisher).

### Animal studies

Our study examined only female mice as ovarian cancer only affects females. All animal studies were performed in accordance with the Institutional Animal Care and Use Committee at the University of Alabama Birmingham. NSG or SCID mice (Jackson Labs) were housed in pathogen-free conditions. For anoikis resistance studies with OV90 cells, SCID (8-week-old NOD.Cg-Prkdcscid/J) mice were injected with OV90-LUC-GFP cells derived from either P0 (parental) or P7 (7 cycles of detachment) that were expanded in TC-treated plates. Live 5×10^6^ cells were injected i.p. and mice were monitored by IVIS imaging every 10 days until day 39-40. Terminal analyses included lung bioluminescence imaging (150µg/ml D-Luciferin), ascites volume, and tumor weight measurements. For ID8 studies, C57BL/6J mice received either parental ID8-EMD cells (P0) or cells after 7 cycles of detachment (P7), expanded in TC-treated plates. Live 10×10^6^ cells were injected i.p. and mice were monitored for 11.5 weeks with weight/girth measurements before endpoint analyses. For SNX631-6 studies, NSG (8week old NOD.Cg-Prkdcscid Il2rgtm1Wjl/SzJ) mice (n=24) received OV90-LUC-GFP cells (5×10^6^, i.p.) and were on either control or SNX631-6 diet (350 ppm, daily dose of 30-50 mg/kg, Senex Biotechnology). These mice were monitored by IVIS imaging and weight/girth measurements every 10 days until day 41, followed by collection of ascites and tumor weight data at necropsy.

### Bulk RNA-seq analysis (secondary analysis and differential gene expression)

Total RNA was isolated using the Trizol/Chloroform extraction method and RNA quality was validated on RNA-1000 chip using Bioanalyzer (Agilent). 1.0 ug of total RNA was used for the construction of sequencing libraries. The RNA-Seq analysis of OV90 P0,P1,P3,P4, P6,P7 samples (Fig. 4) was performed by Novogene, utilizing the NEBNext UltraTM RNA Library Prep Kit for Illumina on an Illumina NovaSeq 6000. All samples contained a minimum of 24 million reads with an average number of 29.2 million reads. The FASTQ files were uploaded to the UAB High-Performance Computer cluster for secondary analysis with the following custom pipeline built in the Snakemake workflow system (v5.9.1)^1^: first, quality and control of the reads were assessed using FastQC, and trimming of the bases with quality scores of less than 20 and adapter were performed with Trim_Galore! (v0.6.4). Following trimming, read quality was re-assessed with FastQC and splice-aware mapping was performed with STAR^2^ (v2.6.0c, with ‘2-pass’ mode) using the GENCODE GRCh38 primary assembly and annotation GTF (release 37). Following genome mapping, BAM index files were generated with SAMtools^3^ (v1.9) and quality control of aligned files was performed with RSeQC^4^ (v3.0.1). Lastly, count generation was performed with ‘featurecounts’ (with ‘Rsubread’^5^, v.1.32.2 and R v3.5.1) and logs of reports were summarized and visualized using MultiQC^6^ (v1.6). The RNA-Seq analysis of OV90 AnS and AnR samples (+/-SNX631, 2D vs 3D, Fig. 8) was performed by University of South Carolina Functional Genomics Core Facility (RRID:SCR_026178), with libraries prepared using NEBNext poly(a) mRNA magnetic isolation kit and NEB Ultra II Directional Library Prep Kit from 500 ng RNA and sequenced on NovaSeq 6000 platform. Processed reads were aligned to genome human GRCh38.p13 primary assembly genome, utilizing STAR (v 2.7.10a) and gene counts were generated by feature counts with the Homo_sapiens.GRCh38.113.gtf annotation file. Tertiary analysis was performed in R (v 4.0.2) with the DESeq2 ^7^ package (v1.34.0). Briefly, pre-filtering of low abundance genes was performed to keep genes that have a mean of at least 5 counts, and normalization was performed. Following count normalization, principal component analysis (PCA) was performed, and genes were defined as differentially expressed genes (DEGs) if they passed a statistical cutoff containing an adjusted p-value <0.05 (Benjamini-Hochberg False Discovery Rate (FDR) method) and if they contained an absolute log_2_ fold change >=1). Additional Pathway analysis was performed using GSVA 1.42.0 [74] utilizing the Hallmarks from the Human MSigDB Collections. The gene signatures for epithelial and mesenchymal gene sets were obtained from. [75] Heatmaps were made with ComplexHeatmap version 2.10.0 [76] with the normalized enrichment score from GSVA using the Euclidean distance method. Hallmark scatterplot graphs use normalized enrichment scores from GSEA with ClusterProfiler and were plotted using GraphPad Prism. The FASTQ files of the current study have been uploaded to NCBI’s Gene Expression Omnibus under accession number GSE241546. Reference for R studio: Posit team (2023). RStudio: Integrated Development Environment for R. Posit Software, PBC, Boston, MA. URL http://www.posit.co/. UpSet plots were made using ComplexUpset R package v 1.3.3. https://cran.r-project.org/web/packages/ComplexUpset/index.html and [77].

### Whole Exome Analysis

Isolation of DNA was performed using Qiagen DNAeasy Blood and Tissue Kit. WES library preparation and sequencing were performed by Novogene utilizing Agilent SureSelect Human All Exon V6 Kit yielding paired-end 150-bp reads on an Illumina NovaSeq 6000 and a median coverage of >159X, for the parental cells P0 and the adapted P7 cells using OV90. Results were provided as demultiplexed paired FASTQs containing the sequencing reads. Sequencing was targeted to a mean depth of at least 100x mean coverage in capture regions. Sequence alignment, quality control, and variant calling were done following GATK best practices using the germline and somatic variant calling algorithms haplotype caller and mutect2, respectively. All sequencing had an alignment of 99.8-99.9%. MAFtools version 2.10.05 [78] was used to compare the sequenced time points. Code Availability: https://github.com/page22emily/RNAseq_Anoikis

### Semi quantitative RT-qPCR

Total RNA was harvested using Trizol/Chloroform extraction. RNA was transcribed using iScript Reverse Transcription Supermix and iTaq Universal SYBR Green Supermix. Expression data was normalized to RPL13A or HPRT or GAPDH. RT-qPCR primer sequences are listed in Table 4.

### Statistics

Experimental data were analyzed using GraphPad Prism (version 10). Data are expressed as Mean ± SEM. *P* < 0.05 and was considered statistically significant. All experiments were performed as at least 3 independent biological trials with multiple technical replicates unless indicated. For comparisons between 2 independent groups, unpaired *t* tests were used. Multiple group comparisons were carried out by the analysis of variance (ANOVA,). For comparisons involving one independent variable, One-way ANOVA followed by Tukey’s multiple comparison was performed, and for two independent variables, Two-way ANOVA followed by Tukey’s multiple comparison was performed.

### Data availability

The RNA sequencing data generated in this study are available at GEO under accession number GSE241546 and GSE309005. Code for data analysis is available at https://github.com/page22emily/RNAseq_Anoikis. Data used to generate figures are available in the supporting data values files. Additional data is available upon request from the corresponding author.

## Authors contribution

MM and RR contributed equally as co–first authors; authorship order among co–first authors was determined by mutual agreement based on contributions and approved by co-authors. KM, MM, AS, and EVB conceived and designed the study. MM, RR, AK, LQM, SS, FM, and MC performed experiments and acquired data. Analysis and interpretation of data were done by KM, MM, RR, EFP, NYL, SS, NH, MKJ, LI, EW, AS, IBR, EVB and MC. Resources and supervision was by KM, NH, AS, IBR, MKJ, EW, and KM. MM, RR, IBR, MC, and KM drafted and edited the manuscript; all authors revised the manuscript critically for important intellectual content and approved the final version. KM obtained funding and administered the project

## ACKNOWLEDGEMENTS

Funding for this work was provided by NIH R01CA230628 to KM and NH, Norma Livingston Ovarian Cancer Foundation to KM, R35GM148351 to AS, R43CA271996 and R01CA266027 to EVB. EVB would like to acknowledge support of the Functional Genomics Core, Microscopy and Flow Cytometry Core, and Drug Design and Synthesis Core of the COBRE Center for Targeted Therapeutics at the University of South Carolina (supported by NIH COBRE grant P20GM109091). KM would like to thank Dr. Amir Jazaeri for a subset of cell lines and would like to acknowledge support from the University of Alabama at Birmingham Biological Data Science Core, RRID:SCR_021766, UAB’s flow cytometry core supported by the Center for AIDS Research, AI027767, The O’Neal Comprehensive Cancer Center, CA013148, UAB Preclinical Imaging Shared Facility supported by P30CA013148 and IVIS S10 instrumentation grant 1S10OD021697, and UAB’s high resolution imaging facility. We also thank UAB Bio-Analytical Redox Biology core facility and Melissa J. Sammy, PhD at the UAB Bio-Analytical Redox Biology (BARB) core, DRC (NIDDK p30DK079626, NORC (NIDDK P30DK056336), CCTS (NIH UL1TR003096). Schematics were made using Biorender.

