## Supplementary Figure for "Anoikis resistance and metastasis of ovarian cancer can be overcome by CDK8/19 Mediator kinase inhibition"

**Supp. Figure 1**

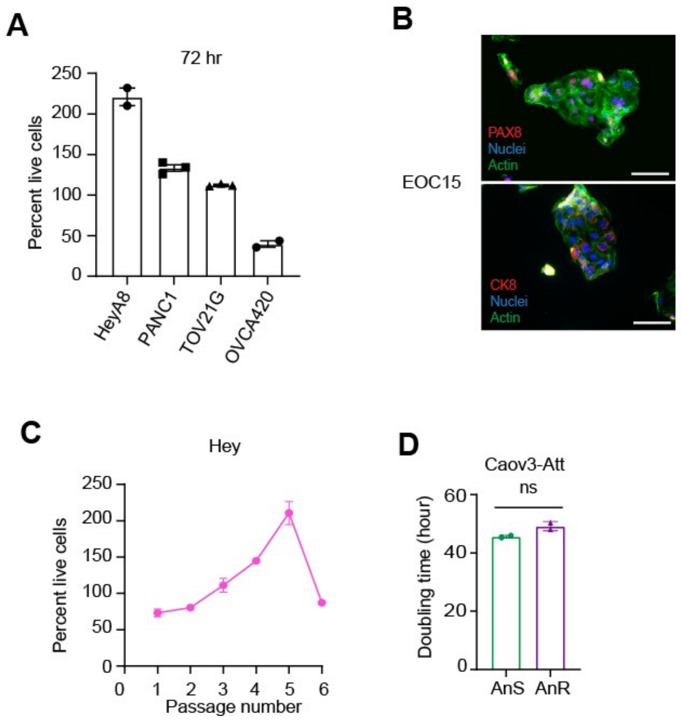


**Supp Figure 1:**

**(A)** Percent live cells in suspension of cell lines HeyA8, PANC1, TOV21G and OVCA420 measured by trypan blue staining following 72 hours incubation in suspension culture (n = 2-3 replicates/cell line). **(B)** Representative immunofluorescence images of F-actin (green), PAX8, and CK8 (red) from EOC15 primary cells, Scale bar: 100 μm. **(C)** Percent live cells in suspension of Hey cells following cyclic model of gain and loss of attachment from Fig 1B. Live cells were measured by trypan blue staining after 24 hours in suspension and plotted as percent relative to initial plating number (n = 2-3 replicates). Live cells were measured by trypan blue staining after 24 hours in suspension and plotted as percent relative to initial plating number (n = 3 replicates) Data are mean ± SEM. ns $p > 0.05$, * $p < 0.05$, ** $p < 0.01$, *** $p < 0.001$, Two-way ANOVA followed by Tukey's multiple comparison. **(D)** Doubling time of parental AnS CAOV3 cells or AnR CAOV3 AnR derivative in attached (Att) growth conditions calculated over a 10-day period using an SRB assay. (Mean ± SEM, ns $p > 0.05$, unpaired t-test.

**Supp. Figure 2**

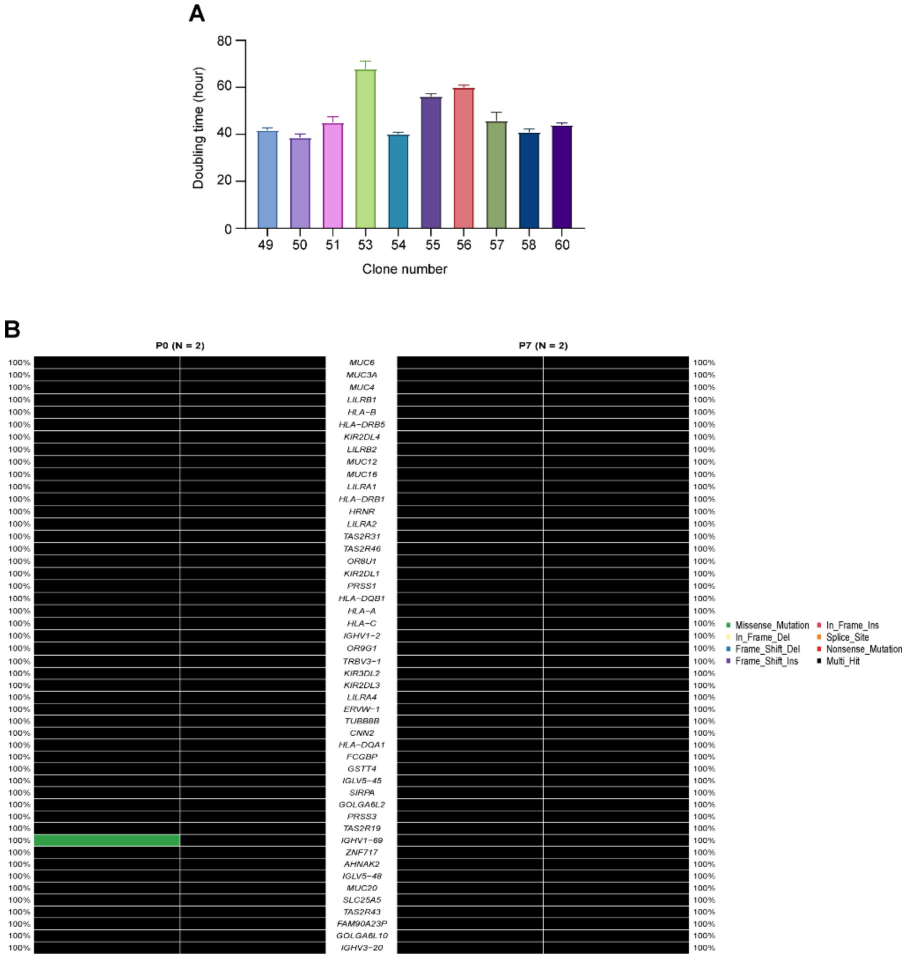


**Supp Figure 2:**

**(A).** Doubling time of indicated clones (randomly chosen) measured by an SRB assay every 24-48 hours over a 7-day period. **(B)**. Oncoplot of the top 50 mutated genes based on the total number of mutations present across 2 time points, P0 (left) and P7 (right).

### Supp. Figure 3

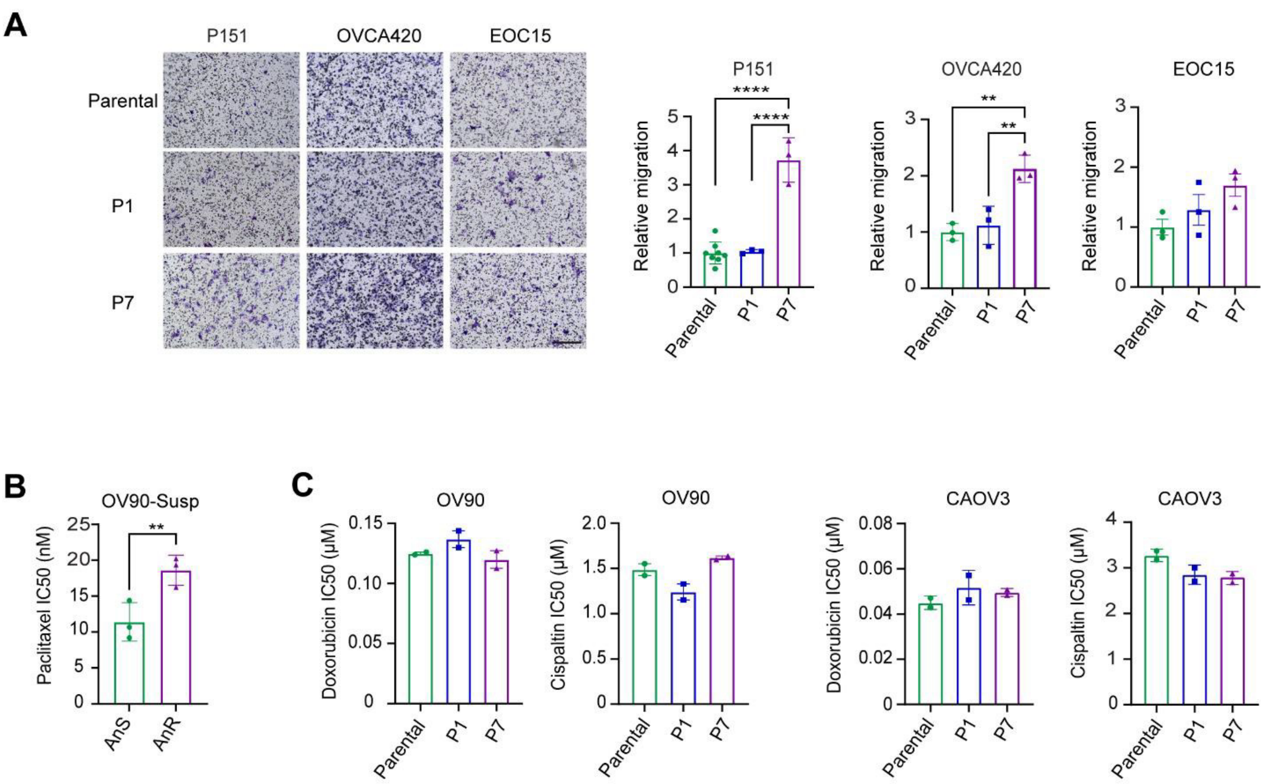


**Supp Figure 3:**

**(A).** Representative images (left) and quantitation (right) of indicated AnS parental, P1 (cells expanded after one 24 hr. exposure to suspension culture) and P7 (AnR) cells on fibronectin coated transwell filters after 24 hrs. of migration (n=3). Data are Mean ± SEM, ns p >0.05,** p < 0.01, ****P<0.0001. One way ANOVA followed by Tukey's multiple comparison. Scale bar: 200µm. **(B).** IC50 of paclitaxel in indicated OV90 cell lines determined under suspension culture conditions over a period of 72 hrs. using CellTiter-Glo 3D cell viability assay. Data are Mean ± SEM, ** p < 0.01, unpaired t test. **(C)** IC50 of indicated parental, P1 (cells expanded after one 24 hr exposure to suspension culture) and P7 (AnR) cells to doxorubicin and cisplatin under steady attached conditions over a period of 72 hrs. using an SRB assay. Data are Mean ± SEM, ns p >0.05. One way ANOVA followed by Tukey's multiple comparison.

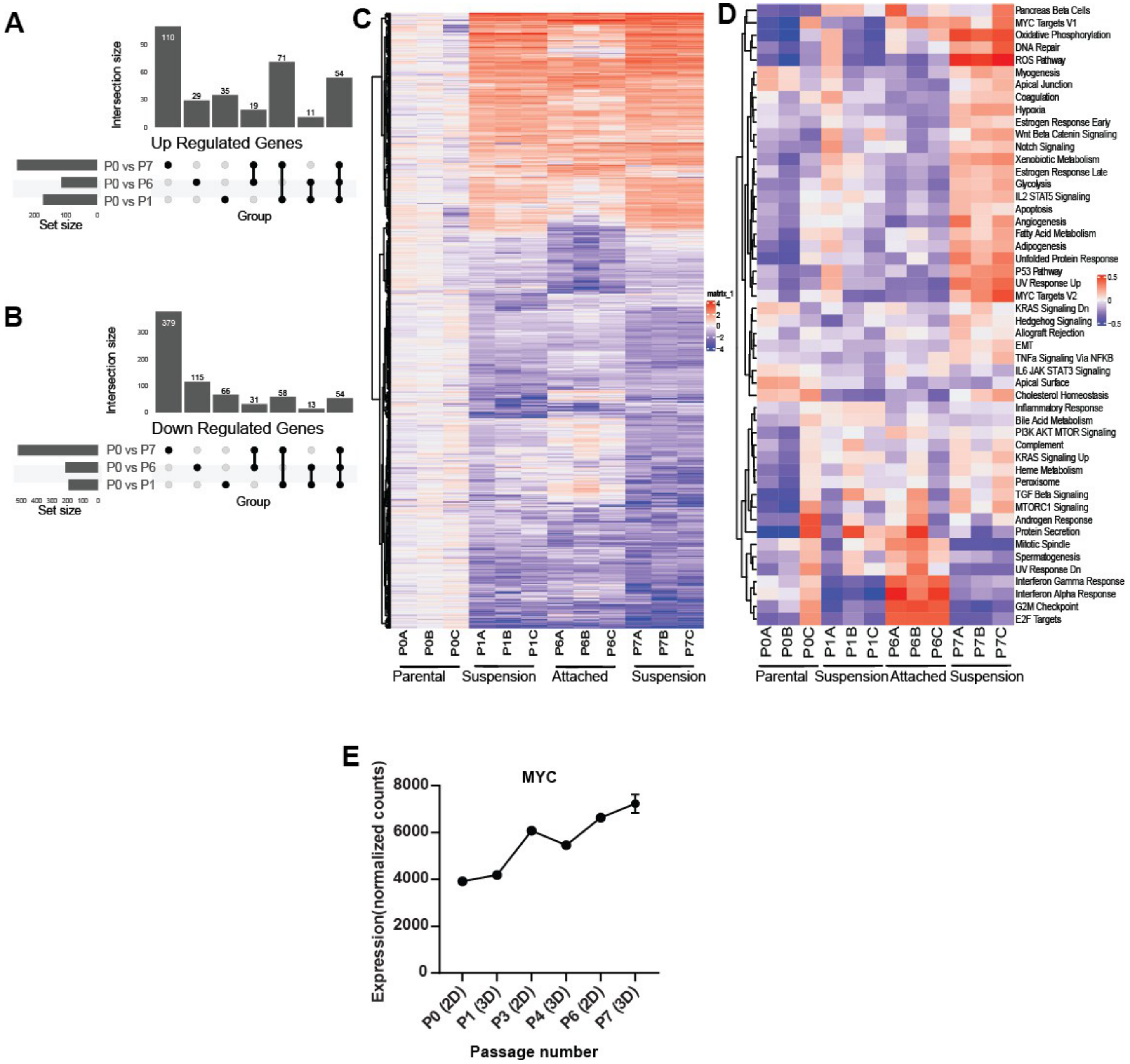


**Supp Figure 4:**

**(A).** UpSet plots indicating the number of up regulated genes (p value < 0.05 and L2FC >1.5) and **(B)** down regulated genes (p value < 0.05 and L2FC < -1.5) in CAOV3 across the time point comparisons P0 vs P1, P0 vs P6, and P0 vs P7. **(C)** Heatmap for the individual CAOV3 samples using Log2FC as calculated for individual biological replicates for parental P0, (P0A-C), P1 (P1A-C), P6 (P6A-C), and P7 (P7A-C). Respective attached and suspension time points are indicated. Heatmap generated by clustering analysis of DEGs across samples (1041 genes). The Log2 fold changes of each gene were clustered based on euclidean distance. **(D).** Heatmaps for the individual CAOV3 samples using GSVA normalized enrichment scores for passage 0 (P0A-C), 1 (P1A-C), 6 (P6A-C), and 7 (P7A-C). The hallmarks were clustered based on euclidean distance. **(E)** c-MYC expression (normalized count from RNA sequencing) in indicated OV90 cells cultured using model of cyclic gain and loss of attachment.

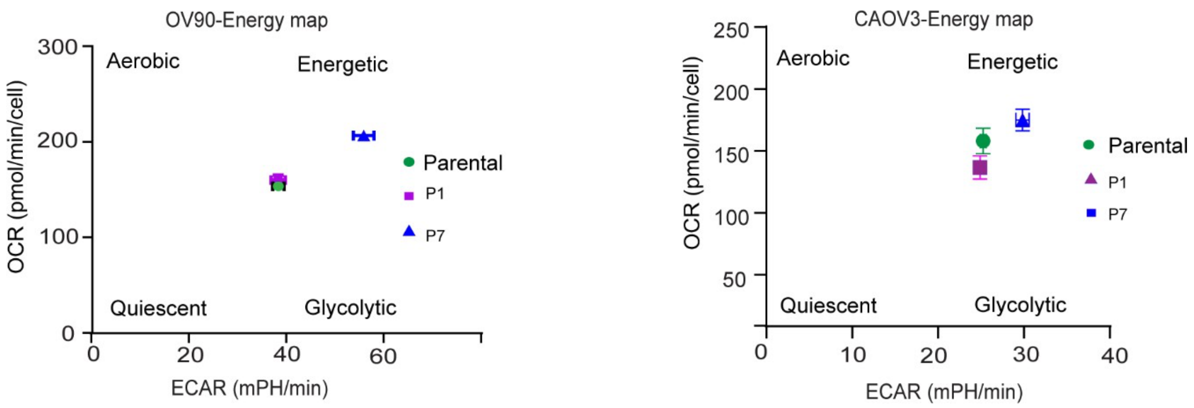


**Supp Figure 5:**

Energy map of indicated OV90 and CAOV3 cells under attached conditions measured using the mito. stress assay on seahorse XFe96 extracellular flux analyzer.

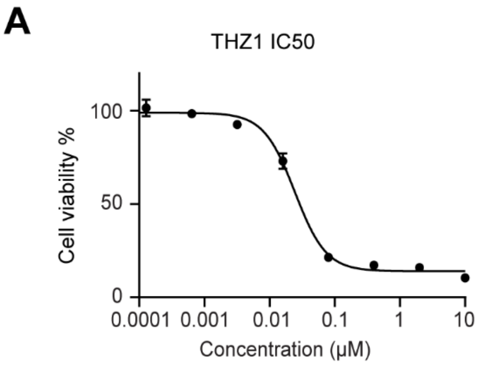


| Cell line | THZ1 Mean IC50 (µM) |
| --- | --- |
| OV90 | 0.034 |

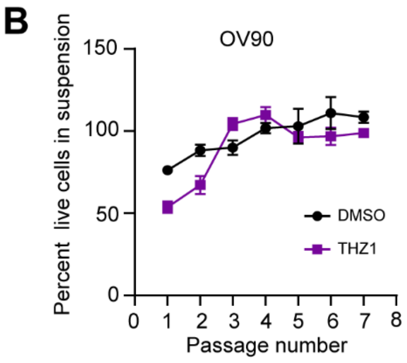


**Supp. Figure 6:**

**(A)** IC50 of OV90 cells assessed after 3 days of incubation with either vehicle or THZ1 using an SRB assay. Adjacent table with calculation of IC50 from. **(B)** Percent survival of OV90 cells measured by trypan blue staining after 24 hours in suspension following cycles of gain and loss of attachment as in Fig 1B, either with vehicle (DMSO) or in the presence of THZ1 (n = 3).
